# Diverse lipid conjugates for functional extra-hepatic siRNA delivery *in vivo*

**DOI:** 10.1101/289439

**Authors:** Annabelle Biscans, Andrew Coles, Reka Haraszti, Dimas Echeverria, Matthew Hassler, Maire Osborn, Anastasia Khvorova

## Abstract

RNAi-based therapeutics show promising clinical data for treatment of liver-associated disorders. However, siRNA delivery into extra-hepatic tissues remains an obstacle, limiting the use of siRNA-based therapies. Here we report on a first example of chemical engineering of lipophilic conjugates to enable extra-hepatic delivery. We synthesized a panel of fifteen lipophilic siRNA and evaluated the impact of their chemical configuration on siRNA tissue distribution profile. Generally, lipophilic conjugates allow siRNA distribution to a wide range of tissues, where the degree of lipophilicity defines the ratio of liver/spleen to kidney distribution. In addition to primary clearance tissues, several conjugates achieve significant siRNA distribution to lung, heart, adrenal glands, fat, muscle. siRNA tissue accumulation leads to productive silencing, shown with two independent targets. siRNA concentrations necessary for productive silencing are tissue and conjugate dependent, varying significantly from 5 to 200 ng/mg. The collection of conjugated siRNA described here enables functional gene modulation *in vivo* in lung, muscle, fat, heart, adrenal glands opening these tissues for future therapeutic intervention.

## Introduction

Therapeutic oligonucleotides—i.e., antisense oligonucleotides, small interfering RNA (siRNA), and aptamers—are emerging as a new class of drugs in addition to small molecules and biologics^1^. Their advantages over conventional drugs include: (i) ease of design, rationally achieved based on sequence information and straightforward screening, leading to drug candidates within short periods of time; (ii) ability to target disease genes previously considered “undruggable”; and (iii) unprecedented potency and duration of effect^2, 3^. Clinical success is dependent on their efficient delivery to disease tissues.

For liver delivery, the trivalent N-acetylgalactosamine (GalNAc) conjugate binds with high specificity and affinity to the asialoglycoprotein receptor on hepatocytes, leading to specific oligonucleotide delivery to and robust gene silencing in hepatocytes^4–7^. This delivery strategy has revolutionized the development of oligonucleotide therapeutics to treat liver diseases, with more than a dozen clinical programs^8^. The success of the GalNAc platform demonstrates that functional tissue delivery of therapeutic oligonucleotides is a foundation for any clinical exploration.

Lipid nanoparticles (LNPs) have been successfully used for siRNA delivery *in vivo*. Most LNP formulations preferentially deliver siRNAs to liver but engineering the lipid composition of LNPs was shown to change their distribution, allowing functional delivery to endothelia, lung, and peritoneal macrophages^9, 10^. Thus, lipid moieties can be used to improve siRNA distribution. Indeed, lipid conjugates have been explored to improve siRNA bioavailable. Cholesterol was the first lipid conjugate developed. After systemic administration, cholesterol-conjugated siRNAs preferentially distribute to liver^11, 12^, but are also distributed to other tissues, including muscle and placenta, where they can induce silencing (Turanov *et al.*, in review)^13^. In addition, local injection of cholesterol-modified siRNAs leads to functional gene silencing in brain, vagina, and skin^14–17^. Other lipid-conjugated siRNAs have been explored, including fatty acids^11^ and vitamins (Vitamin E)^18^ conjugated siRNA, but delivery and efficacy were only evaluated in liver.

Other than the conjugate, the design of the siRNA plays a crucial role in improving siRNA delivery. Indeed, unmodified siRNAs are rapidly degraded and cleared from circulation by kidney filtration, leading to minimal bioavailability in tissues^12^. Thus, full chemical stabilization of siRNAs is essential for conjugate-mediated delivery^19^. The clinical success of GalNAc conjugated siRNAs was only achieved when the siRNAs are extensively modified^20, 21^. Chemical scaffolds that replace every 2’ hydroxyl^22–24^, modify terminal nucleotide linkages^25, 26^, and stabilize the 5’ phosphate^27–30^ maximize *in vivo* activity of siRNAs.

Here we systematically evaluate how lipid conjugates affect siRNA distribution *in vivo*. We synthesized a panel of siRNAs conjugated to fifteen different lipophilic moieties, including saturated and non-saturated fatty acids, steroids, and vitamins, with or without a phosphocholine polar head group. We show that lipid conjugates enable siRNA accumulation and productive silencing in a wide range of tissues in mice after subcutaneous injection. In general, the degree of siRNA lipophilicity correlates with accumulation in kidneys (less lipophilic) or liver (more lipophilic). Though most of the injected siRNA accumulates in clearance organs, we identify conjugates that enable functional siRNA delivery to heart, lung, fat, muscle, and adrenal gland. The tissue concentrations of siRNA required for productive silencing varied depending on the tissue and lipid conjugate. Our findings provide proof of principle for the development of lipid-conjugated oligonucleotides as therapeutics for multiple indications.

## Results

### Synthesis of conjugated siRNAs library

To evaluate how the structures of lipid conjugates affect siRNA distribution *in vivo*, we synthesized a panel of siRNAs conjugated with naturally occurring lipids, including: the unsaturated fatty acids 22:6 n-3, docosahexaenoic acid (DHA) and 20:5 n-3, eicosapentaenoic acid (EPA); the saturated fatty acid 22:0, docosanoic acid (DCA); the sterols cholesterol (chol) and lithocholic acid (LA); and the vitamins retinoic acid (RA) and α-tocopheryl succinate (TS) (**Figure 1**). A significant fraction of circulating fatty acids are esterified with phosphatidylcholine or other head groups^31, 32^. We therefore synthesized each lipid conjugate with or without a phosphocholine (PC) head group^33^, to test whether the polar head group affects the distribution profile of lipid-conjugated siRNAs *in vivo*. We also synthesized unconjugated siRNAs (Unconj.) and siRNAs with the PC head group (PC) but no lipid for controls. Each lipid conjugate was covalently attached to the 3’ end of the siRNA sense strand, which tolerates a range of covalent modifications^12, 34, 35^. The siRNAs used in these studies were fully chemically modified for maximal stability and minimal innate immune activation as described (see **Figure 1a** and **Methods**). We and others have recently shown that full chemical stabilization is essential for conjugated-mediated delivery^19–21^.

**Figure 1.**
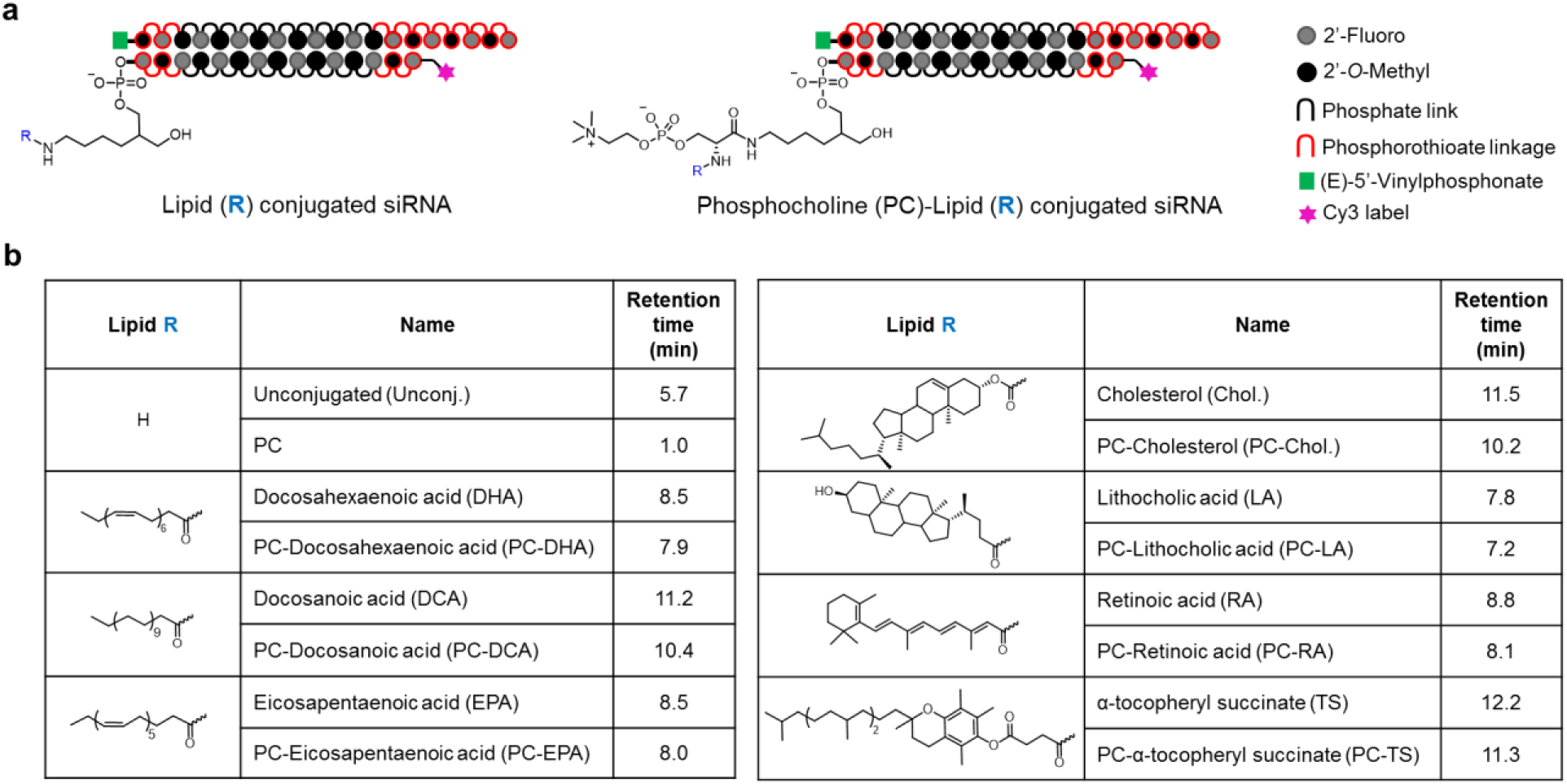
Lipid conjugated siRNAs have different lipophilicity depending of the structure of the lipid. (a) Schematic structures of lipid-conjugated siRNAs. The lipid conjugate (R) is attached to the 3’ end of the sense strand of the siRNA though a C7 linker or a C7 linker functionalized with phosphocholine (PC). (b) Lipid moieties attached to siRNAs and reversed-phase HPLC retention times of conjugated sense strands. Retention time directly relates to hydrophobicity of the oligonucleotide.

The chemical compositions of lipids had a profound effect on the hydrophobicity of lipid-siRNA, based on retention time in reversed-phase HPLC^36^, which ranged from 1 to 12.2 minutes (**Figure 1b**). In general, compounds with a PC head group eluted 0.5 to 1.3 min earlier than counterparts without a PC head group, suggesting that the PC head group reduces lipophilicity. Oligonucleotides without a lipid eluted the fastest, with the PC-siRNA conjugate (1 min) eluting 4.7 min earlier than unconjugated oligonucleotide control (5.7 min). Lipid-conjugated siRNAs fell into two categories: low lipophilic compounds (retention times from 7.2 to 8.8 min) with unsaturated carbon chains or cyclic structures (LA, PC-LA, EPA, PC-EPA, RA, PC-RA, DHA, and PC-DHA conjugates); and high lipophilicity compounds (retention times from 10.2 to 12.2 min) with saturated carbon chains (TS, PC-TS, DCA, PC-DCA, chol. and PC-chol. conjugates).

### Lipid conjugates define *in vivo* distribution of lipophilic siRNAs

To evaluate how the lipid moiety affects siRNA distribution to tissues, mice were injected subcutaneously with lipid-conjugated siRNA or PC-siRNA or unconjugated siRNA (3 mice per compound). The siRNA sense strands were labeled at their 5’-ends with a Cy3 fluorophore for qualitative analysis of tissue sections by fluorescence microscopy. We also quantified the concentrations of siRNA antisense strands in tissue biopsies using a peptide nucleic acid (PNA) hybridization assay^37, 38^. It is important to note that the PNA hybridization assay is not dependent on the presence of a Cy3 dye. Indeed, the levels of tissue accumulation of Cy3-labeled and non-labeled siRNA were compared for several conjugated siRNAs and were shown to be similar (data not shown). In addition, we have recently shown that conjugated siRNAs are cleared from the blood stream and accumulate in liver and kidneys within hours of injection, and the tissue distribution of siRNA 48 hours after injection is predictive of the long-term retention of siRNA^38^. We therefore analyzed the distribution of lipid-conjugated siRNAs, 48 hours after injection, in 15 tissues: liver, kidneys, adrenal glands, lung, heart, thymus, spleen, pancreas, intestine, fallopian tube, bladder, fat, muscle, injection site, and skin. **Supplementary Figures 1-15** shows fluorescence microscopy images (5× and 40× magnification) of each tissue for each lipid-siRNA conjugate, and quantification of antisense strand concentration in each tissue. The quantitative data for all 16 compounds in all 15 tissues are summarized in **Figure 2a** using a heat map.

**Figure 2.**
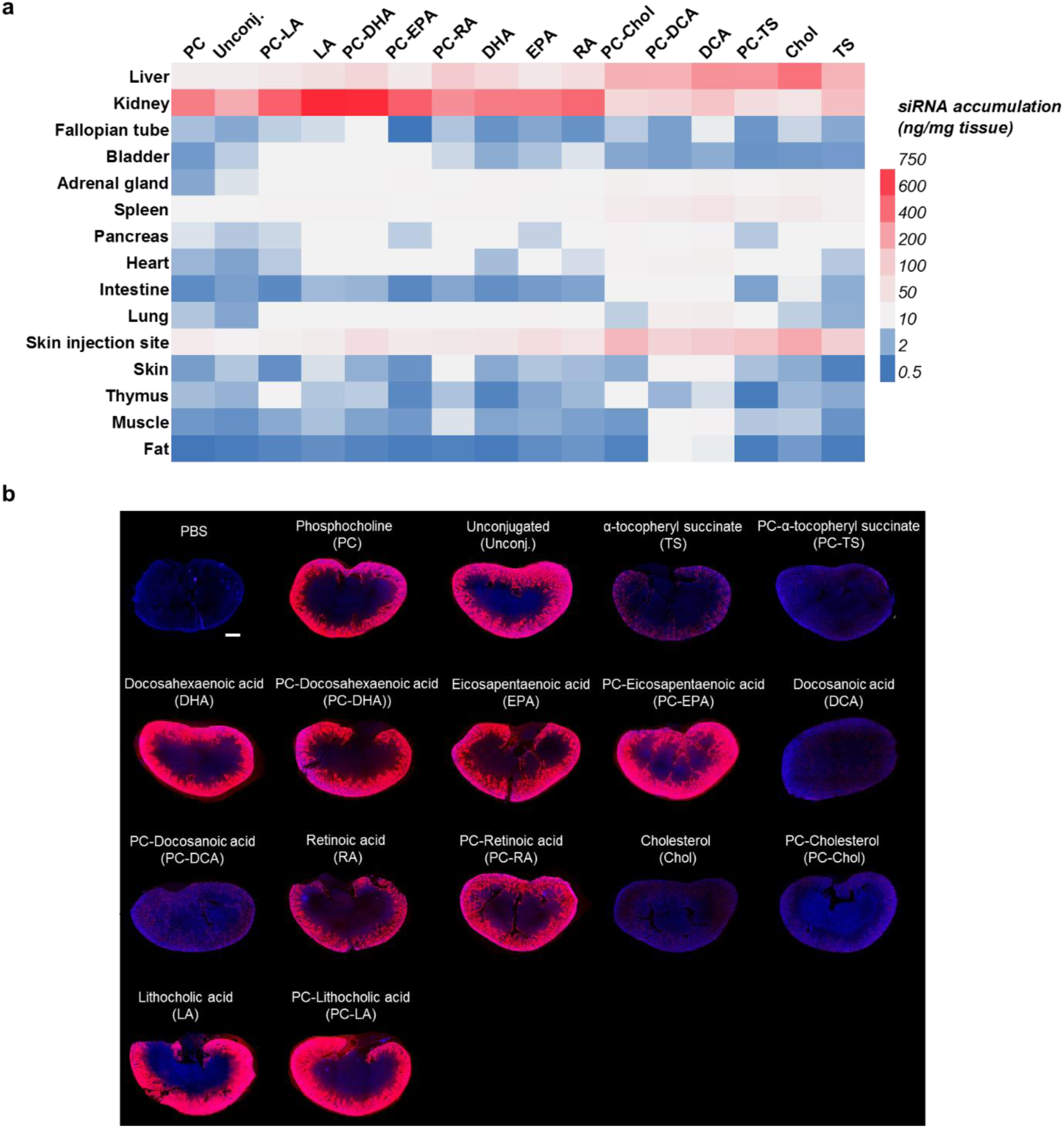
The structure of the conjugate defines siRNA distribution. (a) Heat map representation of the tissue concentrations of siRNA antisense strands. Data represent the average of 3 experiments. (b) Representative fluorescence images of kidney sections from mice injected subcutaneously with 20 mg/kg Cy3-labeled lipid-conjugated siRNAs (red). Nuclei stained with DAPI (blue). Three mice injected per conjugate. Images taken at 5× magnification and collected at the same laser intensity and acquisition time. Scale, 1 mm.

As expected, conjugated siRNAs accumulated in primary clearance tissues (kidneys and liver), at the injection site, and in other tissues with discontinuous or fenestrated epithelia (e.g., spleen). We observed a clear relationship between siRNA lipophilicity and the amount of siRNA that accumulated in these tissues. For example, low lipophilic siRNAs (DHA-, EPA-, RA-, and LA-siRNAs, and PC-conjugated derivatives) accumulated to higher levels in kidneys than in the liver or spleen or at the injection site (**Figure 2, Supplementary Figures 1, 2, 3 and 14**). By contrast, high lipophilic siRNAs (TS-, DCA-, and Chol-siRNAs, and PC-conjugated derivatives) showed limited kidney distribution (**Figure 2b**) and accumulated to higher levels in liver and spleen and at the site of injection than low lipophilic siRNAs (**Figure 2a**). Unconjugated siRNAs also accumulated to high levels in kidneys, due to the presence of phosphorothioate linkages^25, 39, 40^. The retention of high lipophilic siRNAs at the injection site agrees with a recent study comparing the clearance kinetics of PC-DHA- and cholesterol-conjugated siRNAs^38^.

Although the highest levels of siRNA were detected in liver and kidneys, significant levels of siRNA were detected in other major tissues, including heart (∼20 ng/mg), lung (∼29 ng/mg), muscle (∼9 ng/mg), fat (∼4 ng/mg), bladder (∼5 ng/mg), spleen (∼56 ng/mg), and adrenal glands (∼20 ng/mg) (**Supplementary Figures 3-7, 11, 13**). In most tissues, high lipophilic siRNAs accumulated to the highest levels. Thus, lipid moities allowed siRNA delivery to tissues throughout the body.

Lipid structure also affected siRNA distribution within tissues. In kidneys, siRNAs primarily accumulated in proximal epithelia, consistent with its role in filtration (**Supplementary Figure 2**). Nevertheless, some lipid-conjugated siRNAs (e.g. PC-RA- or EPA-siRNAs) accumulated to low but detectable levels in medulla, glomerulus and bowel capsule. The effect of lipid conjugate was even more striking in liver (**Supplementary Figure 1**). For example, cholesterol-conjugated siRNA variants were detected in all liver cell types, including Kupffer cells, endothelial cells, and hepatocytes. DCA-conjugated siRNAs were also detected in all three cell types, but its levels in Kupffer and endothelial cells appeared to be higher than in hepatocytes. Low lipophilic conjugates, e.g. DHA, were mostly detected in endothelial cells.

The tissues that we analyzed comprise most of the mouse body, and the relative mass of each tissue has been experimentally defined or estimated^41–43^. Our ability to quantify the concentration of siRNAs in tissues therefore allowed us to estimate the fraction of the injected dose of siRNA retained 48 hours after injection and the fraction of retained siRNA per tissue (**Figure 3**). Based on these calculations, we estimated near complete retention (>70%) of high lipophilic compounds (**Figure 3a**), because only a small fraction of the injected dose was accumulated to and cleared though kidneys (**Figure 3b**). By contrast, for most low lipophilic compounds, mice retained less than 60% of the injected dose, and most of the retained dose resided in the kidneys (**Figure 3**), showing important renal clearance. Only 15% of unconjugated siRNAs were retained, and ∼60% of the retained dose resided in kidneys (**Figure 3**). This analysis revealed one notable outlier: mice retained <50% of the high lipophilicity PC-Chol-siRNA, with ∼70% of the retained dose residing in liver.

**Figure 3.**
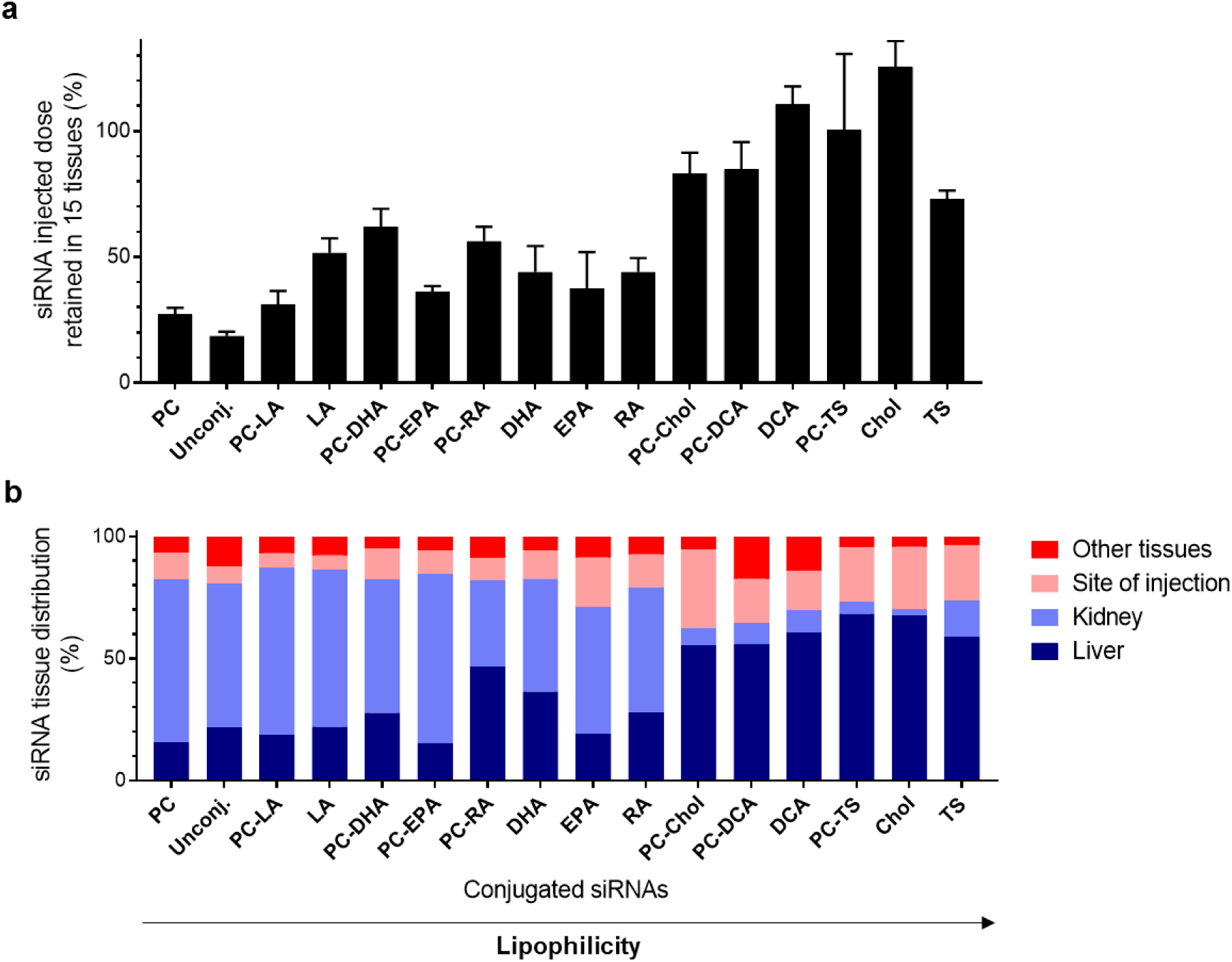
The structure and lipophilicity of the conjugate predict siRNA distribution. (a) Bar graph showing the percent of the injected dose retained in mice across all 15 tissues (n=3). (a) Bar chart showing the percent of retained siRNAs that reside in liver, kidney, site of injection (skin), and all other tissues. siRNAs are sorted by increasing relative lipophilicity.

Aside from reducing the lipophilicity of lipid-conjugated siRNAs, the PC polar head group had variable effects on the clearance and tissue accumulation of siRNAs. The addition of a PC moiety increased the overall retention of TS-, DHA-, and RA-conjugated siRNAs, but increased the clearance of DCA-, Chol-, and LA-conjugated siRNAs (**Figure 3a**). Nevertheless, the addition of a PC head group appeared to improve the retention of RA-, Chol- and DCA-conjugated siRNAs in tissues other than liver, site of injection and kidneys (**Figure 3b**). Indeed, the PC-DCA-siRNA conjugate showed the highest level of retention (17% of retained dose) in non-clearance tissues. These results suggest that although clearance of siRNA is inversely related to lipophilicity, some conjugates regulate clearance and distribution by as yet unknown mechanisms.

### Lipid-conjugated siRNAs enable functional gene silencing in several extra-hepatic tissues, including heart, lung, muscle, fat, adrenal glands, and kidneys

To determine if lipid-conjugated siRNAs accumulate to levels sufficient for productive silencing, we injected mice subcutaneously with lipid-conjugated siRNAs targeting *Huntingtin* (*Htt*)^17^ or *Cyclophilin B* (*Ppib*)^44^ mRNAs. We chose these targets because validated siRNA sequences are available, both targets are widely expressed in the body, and both targets are expressed at different levels (*Htt* low, and *Ppib* high). We injected each conjugated siRNA at a dose of 20 mg/kg (16 mice per conjugate including non-targeting controls), and a week later we measured *Ppib*, *Htt*, and *Hprt* (hypoxanthine-guanine phosphoribosyl transferase, serving as a housekeeping gene) mRNA levels. At the dose tested (20 mg/kg), all conjugated siRNAs were well tolerated: we observed no adverse events or changes in blood chemistry (**Supplementary Figure 31**). For tissues that retained, for the majority of the compounds, more than 5 ng of conjugated siRNA per mg of tissue (liver, kidney, skin injection site, spleen, adrenal glands, heart, lung, and skin), we measured the activity of all 15 conjugated siRNAs and the unconjugated siRNA. For the remaining organs (intestine, bladder, fat, muscle, pancreas, thymus, and fallopian tube), we only measured the activity of conjugated siRNAs that accumulated to the highest level, level dependent on the tissue. In total, we processed around 3,000 samples. The silencing efficiency of each compound in each tissue is shown in **Supplementary Figures 16 to 30**, and data are summarized as heat maps in **Figure 4**. In all cases, non-targeting controls (*Ntc*) showed no significant reduction in target gene expression, indicating that the observed silencing is due to sequence specific effects rather than the general chemical scaffold.

**Figure 4.**
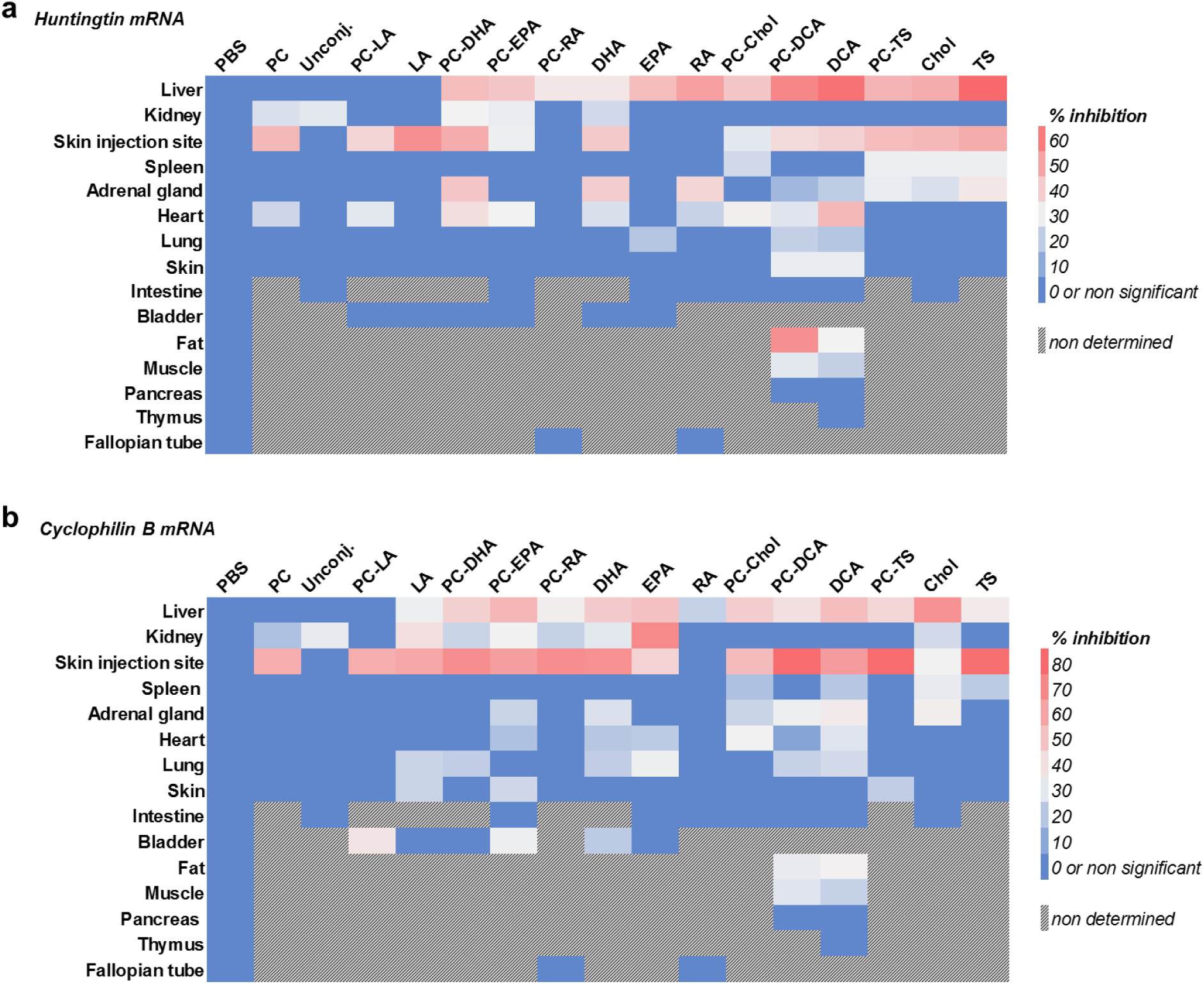
mRNA silencing depends on the lipid conjugate attached to the siRNA. (a, b) Heat map representations summarizing the levels of silencing of *Huntingtin* mRNA (a) or Cyclophilin B mRNA (b) by each siRNA in each tissue as a percent of *Huntingtin* and *Cyclophilin B* mRNA levels in PBS control-treated mice. Each data point represents the mean of 8 injected mice.

Unconjugated siRNAs only induced silencing in kidneys, demonstrating that statistically significant silencing depends on conjugate-mediated delivery (**Figure 4, Supplementary Figures 16-30**). With one exception, all of the conjugated siRNAs induced statistically significant downregulation of *Htt* or *Ppib* at the injection site (**Figure 4, Supplementary Figure 22**). Despite a high level of retention, RA-siRNA was the only conjugate that failed to silence either target at the injection site. Similarly, most of the conjugated compounds (except PC-, and PC-LA-siRNAs) induced silencing of *Htt* or *Ppib* in the liver (**Figure 4, Supplementary Figure 16**). In addition to liver and injection site (skin), in general, high lipophilic siRNAs induced significant levels of silencing in spleen (24% to 30% for *Htt*; 18% to 31% for *Ppib*) and adrenal glands (20% to 34% for *Htt*; 24% to 36% for *Ppib*) (**Figure 4, Supplementary Figures 18, 21**). Low lipophilic siRNAs showed significant silencing activity in kidneys (**Figure 4, Supplementary Figure 17**).

To examine the relationship between siRNA accumulation and efficacy in tissues, we plotted the level of silencing against the tissue concentration for each compound (**Figure 5**).

**Figure 5.**
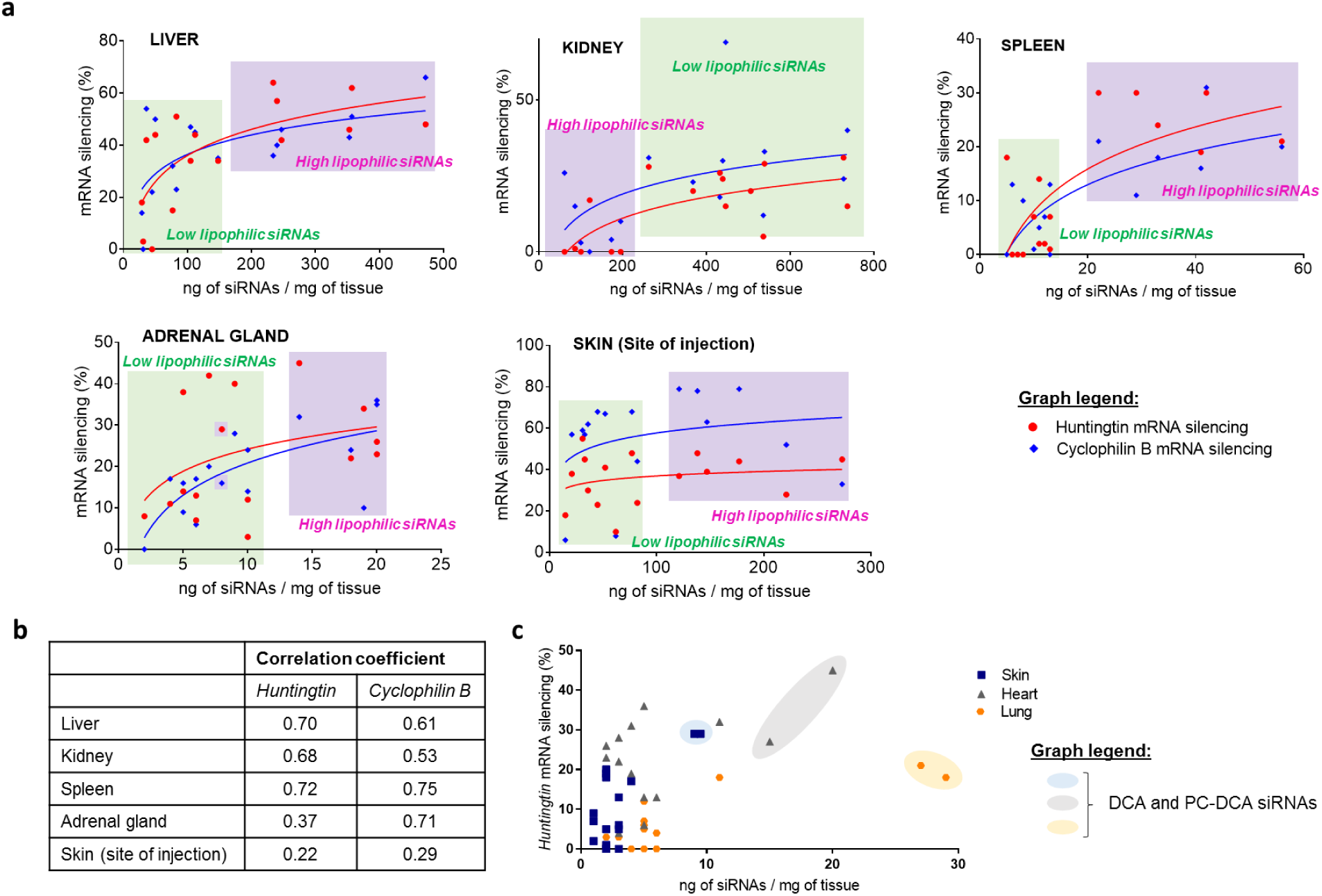
The degree of mRNA silencing correlates with the siRNA distribution in tissues. (a) Dot plots showing percent silencing of *Huntingtin* or *Cyclophilin B* mRNAs by lipid-conjugated siRNAs plotted against tissue concentration of the siRNA in liver, kidney, spleen, adrenal gland, and skin (site of injection). Low and high lipophilic siRNAs tend to cluster together. Low lipophilic siRNAs (green shading) include PC, Unconj., PC-LA, LA, PC-DHA, PC-EPA, PC-RA, DHA, EPA, and RA conjugates. High lipophilic siRNAs (purple shading) include PC-Chol, PC-DCA, DCA, PC-TS, Chol, and TS conjugates. Lines represent best fit of data. (b) Correlation coefficients showing the strength of the relationship between silencing and siRNA accumulation from data in (a). (c) Dot plot of percent Huntingtin silencing in skin, heart, and lung by lipid-conjugated siRNAs plotted against tissue concentration of the siRNA. The data indicate that DCA- and PC-DCA-conjugated siRNAs to higher levels and silence *Huntingtin* better than other conjugated siRNAs.

This analysis again clearly shows relationships between lipophilicity and tissue accumulation of conjugated siRNAs, as low and high lipophilic siRNAs tended to cluster separately. We also observed a moderate positive correlation between siRNA accumulation and silencing activity in several tissues (r = 0.53 to 0.75, **Figure 5b**), but the correlation did not always hold for both targets. Moreover, different compounds sometimes showed dramatically different activities, even if they were present at similar concentrations. In liver, for example, PC- and PC-EPA-siRNAs accumulated to similar levels (31 ng/mg vs. 36 ng/mg). However, whereas PC-EPA-siRNA induced ∼50% silencing, PC-siRNA was completely inactive. Conversely, liver accumulation of DCA-siRNA was 10-fold higher (357 ng/mg) than that of PC-EPA-siRNA, but both compounds induced similar levels of silencing. A similar trend was observed in other tissues (**Figure 5a**). These findings suggest that the structure of the lipid conjugate can have a profound effect on the biological activity of siRNAs within a tissue.

The threshold of lipid-conjugated siRNA required to down-regulate *Ppib* or *Htt* was also tissue-dependent. Whereas tissue concentrations of siRNA greater than 250 ng/mg were required for silencing in kidney tissue, concentrations between 20 and 60 ng/mg induced silencing in spleen (**Figure 5a**). Even lower concentrations of siRNA were sufficient to induce potent silencing in fat (4 ng/mg) and muscle (7 to 9 ng/mg) (**Supplementary Figures 11, 13, 25-26**). This analysis also showed that *Htt* and *Ppib* were more or less sensitive to silencing depending on the tissue. For example, *Ppib* was more effectively silenced than *Htt* at the injection site, whereas *Htt* was more effectively silenced than *Ppib* in adrenal glands, likely due to differences in relative levels of gene expression in the targeted cellular population (**Figure 5a**). Thus, in addition to the structure of the compound, the biology of the tissue affects the mechanism of uptake and biological availability.

These results suggest that (i) the amount of siRNAs required to induce silencing is tissue dependent, (ii) in some tissues a large amount of siRNA is not necessary to induce silencing, and (iii) over a certain amount of siRNAs in the tissue, a portion of the siRNAs is trapped in a non-productive pathway^45, 46^.

### DCA and PC-DCA-conjugated siRNAs enable wide-spread silencing

Several compounds, particularly DCA and PC-DCA-conjugated siRNAs, accumulated to significant levels in tissues other than liver and kidney (**Figure 5c, Figures 6a-6d**), where they induced productive silencing (**Figure 5c, Figures 6e-6f**). The degree of silencing depended on the target, but both DCA-and PC-DCA-conjugated siRNAs were able to significantly silence *Htt* or *Ppib* in heart, lung, adrenal glands, muscle, and fat. Depending on the tissue, the PC group either reduced, increased or unchanged *Htt* or *Ppib* silencing.

**Figure 6.**
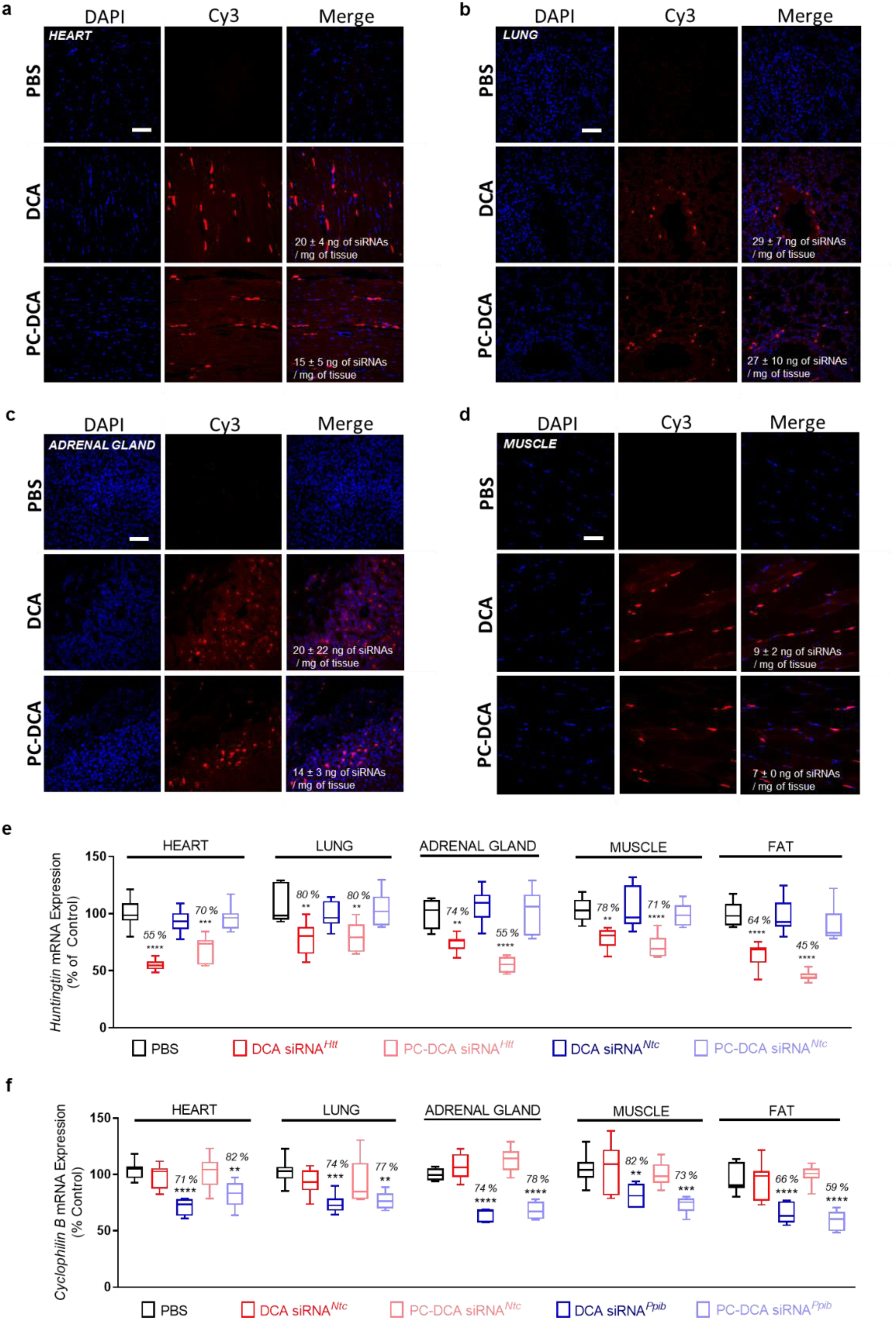
DCA and PC-DCA conjugated siRNAs are the best candidates to induce silencing in tissues. (a-d) Fluorescence images of Cy3-labeled DCA- and PC-DCA-conjugated siRNAs in (a) heart, (b) lung, (c) adrenal gland, and (d) muscle. Scale, 50 µm (e) *Huntingtin* and (f) *Cyclophilin B* mRNA levels in heart, lung, adrenal gland, muscle, and fat. In each tissue, *Huntingtin* and *Cyclophilin B* mRNA levels were normalized to *Hprt* mRNA levels. Non-targeting control (*Ntc*) siRNAs target *Cyclophilin B* in (e) and Huntingtin in (f). Data are shown as percent of PBS control. ****P<0.0001, ***P<0.001, **P<0.01.

Taken together, our findings indicate that a single subcutaneous injection of DCA- or PC-DCA-conjugated siRNAs supports functional target regulation in multiple tissues throughout the body, opening these tissues to RNAi-based functional genomics studies.

## Discussion

GalNAc-conjugated siRNAs have dominated the recent development of therapeutic oligonucleotides for liver indications. The GalNAc moiety drives specific delivery to and activity in hepatocytes^4–6^. We have shown that lipid conjugates support much broader delivery of siRNAs and enable functional silencing in many tissues, including liver, kidneys, lung, heart, muscle, spleen, fat, and adrenal glands. To our knowledge, only one other study has reported siRNA delivery to muscle after systemic injection^13^. At the dose tested, we did not observe overt toxicity or abnormal blood chemistry. Our findings suggest that, with careful selection, lipid conjugates promise to advance the therapeutic potential of siRNA.

For siRNAs to be delivered throughout the body, they must be retained. Lipid conjugates were essential for retention of siRNA in the body. Indeed, ∼85% of unconjugated siRNA was rapidly cleared from the body, and most of the retained siRNA accumulated in kidneys. Unconjugated siRNAs are ∼13 kDa—well within the limit (approximately 40 kDa) filtered by the kidney. Clearance is affected by the nature of the conjugate and is slower for high lipophilic compounds^38^. Here we estimate that whereas retention of low lipophilic compounds varied from 27% to 62% of the injected compound, high lipophilic (e.g., cholesterol, DCA, and α-tocopheryl succinate) siRNA conjugates are almost completely retained by the body. The ability of lipophilic conjugates to bind to serum proteins is known to define *in vivo* distribution of small molecules. Indeed, lipid conjugates have a profound impact on siRNA binding to serum proteins, which correlates with observed tissue distribution pattern^11^ (Osborn *et al.*, 2018, in preparation). Future studies are needed to identify additional mechanisms that contribute to the *in vivo* retention and distribution of siRNAs.

The potential for widespread delivery of siRNAs comes with shortcomings. Lipid-conjugated siRNAs are not delivered to specific tissues, and most of the injected dose will be delivered to primary clearance tissues, including liver, kidney, and spleen. Nevertheless, lipid-siRNA conjugates could be used to silence relevant targets that are only expressed in the disease tissue, or to silence disease targets that are widely expressed as long as the loss of activity is tolerated in healthy tissues. For example, fully chemically stabilized chol-conjugated siRNAs have been used to silence soluble FTL1, which is overexpressed in placentas of pregnant subjects with preeclampsia (Turanov *et al*., 2018, under review). Though the chol-siRNA targeting sFLT1 is efficiently delivered to placenta (∼8% of injected dose), it also accumulates to liver and kidneys where silencing of sFLT1 is believed to be tolerated and irrelevant for disease progression. There are targets of the similar nature, including DUX4 in muscular dystrophy^47^. Thus, use of lipophilic conjugates offers a potential for modulation of gene expression in many tissues, as long as target and clinical indication are carefully considered and matched to the pharmacologic and safety profile of the lipid-siRNA conjugate.

Lipid-based nanoparticles (LNP) have been explored as a way to deliver siRNA for decades. Years of chemical evolution have substantially improved LNPs efficacy and safety^9, 48^. The size of an LNP particle has a dramatic effect on distribution, with liver being a predominant target tissue^49^. Indeed, differences in fenestrated capillary between tissues lead to the extravasation of particles with different diameters: ∼2 nm in heart, muscle, lung, and skin; ∼30 nm in kidneys; and ∼150 nm in liver and spleen. The direct attachment of lipids to siRNAs supports delivery to a much wider array of tissues, because lipid-siRNA conjugates behaves more like small molecules bound to the serum proteins.

This study reports a first example of chemically engineered lipid-conjugated siRNAs for systemic delivery. Future expansion of the chemical space of lipophilic moieties will establish a path towards enhancing siRNA delivery and potency.

## Methods

### Synthesis of lipid functionalized solid support for the preparation of conjugated siRNAs

Lipid moieties (except α-tocopheryl succinate) were directly attached via a peptide bond to a controlled pore glass (CPG) functionalized by a C7 linker, as described^50^. To synthesize phosphocholine derivatives, amino C7 CPG was first functionalized with phosphocholine essentially as described^33^. Briefly, Fmoc-L-serine tert-butyl (TCI America) was phosphitylated using 2’-cyanoethyl-N,N-diisopropylchlorophosphoramidite (ChemGenes). The resulting phosphoramidite was coupled to choline p-toluenesulfonate (Alfa Aesar) using 5-(ethylthio)-1H-tetrazole (ETT) as an activator. The phosphine ester was then oxidized, and the carboxylic acid and phosphate ester groups were deprotected (i.e., tert-butyl and cyanoethyl groups removed). The resulting intermediate was attached to the amino C7 CPG via a peptide bond to form phosphocholine-functionalized CPG. The Fmoc group was then removed, and the selected lipid moiety was attached via a peptide bond to the CPG. All lipid-functionalized solid supports were obtained with a loading of 55 μmol/g.

### Synthesis of α-tocopheryl succinate-conjugated siRNAs

α-tocopheryl succinate was attached to the amino group at the 3’ ends of siRNA sense strands after synthesis, deprotection and purification of sense strands synthesized on amino C7 CPG or phosphocholine-functionalized amino C7 CPG. *N*-hydroxysuccinimide α-tocopheryl succinate and purified sense strands were combined in a solution of 0.1 M sodium bicarbonate, 20% (v/v) dimethylformamide and incubated overnight at room temperature. One-tenth volume 3 M sodium acetate (pH 5.2) was added to obtain a final concentration of 0.3 M sodium acetate. Three volumes 95% (v/v) ethanol were added, and the mixture was vortexed and then placed for 1h at −80°C. The solution was pelleted by centrifugation for 30 minutes at 5200 × *g*. The pellet containing the lipid-conjugated siRNA sense strand was dissolved in water, purified, and desalted as described below.

### Oligonucleotide synthesis

An Expedite ABI DNA/RNA synthesizer was used to synthesize oligonucleotides following standard protocols. Sense strands were synthesized at 10 µmole scales on lipid-functionalized CPG or amino C7 CPG (for the post-synthetic conjugation of α-tocopheryl succinate moiety) supports. Antisense strands were synthesized at 10 µmole scales on CPG functionalized with Unylinker^®^ (ChemGenes, Wilmington, MA). 2’-*O*-methyl phosphoramidites (ChemGenes, Wilmington, MA), 2’-fluoro phosphoramidites (BioAutomation, Irving, Texas), Cy3-labeled phosphoramidites (Gene Pharma, Shanghai, China), and custom synthesized (E)-vinylphosphonate phosphoramidites^33^ were prepared as 0.15 M solutions in dry acetonitrile. Trityl groups were removed using 3% dichloroacetic acid (DCA) in dichloromethane for 80 s, phosphoramidites were coupled using 0.25 M 5-(benzylthio)-1H-tetrazole (BTT) in acetonitrile as an activator for 250 s, and uncoupled monomers were capped with 16% N-methylimidazole in tetrahydrofuran (CAP A) and 80:10:10 (v/v/v) tetrahydrofuran:acetic anhydride:2,6-lutidine (CAP B) for 15 s. Sulfurizations were carried out with 0.1 M solution of 3-[(dimethylaminomethylene)amino]-3H-1,2,4-dithiazole-5-thione (DDTT) in acetonitrile for 3 min. Oxidation was performed in 0.02 M iodine in tetrahydrofuran:pyridine:water (70:20:10, v/v/v) for 80 s.

### Deprotection and purification of oligonucleotides

Sense strands were cleaved and deprotected using 1 ml of 40% aq. methylamine at 45 °C for 1h. Antisense strands were first deprotected with a solution of bromotrimethylsilane:pyridine (3:2, v/v) in dichloromethane (5 ml) for the (E)-vinylphosphonate deprotection and then cleaved and deprotected with 10 ml of 40% aq. methylamine at 45 °C for 1h. Sense and antisense oligonucleotide solutions were frozen in liquid nitrogen for a few minutes and dried overnight in a Speedvac under vacuum. The resulting pellets were suspended in water and purified using an Agilent Prostar System (Agilent, Santa Clara, CA). Sense strands were purified over a Hamilton HxSil C18 column (150 × 21.2) in a continuous gradient of sodium acetate: 90% Buffer A1 (50 mM sodium acetate in 5% acetonitrile), 10% Buffer B1 (acetonitrile) to 10% Buffer A1, 90% Buffer B1 at a flow rate of 5 ml/min for 18 min at 70°C. Antisense strands were purified over a Dionex NucleoPac PA-100 (9 × 250) in a continuous gradient of sodium percholate: 100% Buffer A2 (30% acetonitrile in water) to 20% Buffer A2, 80% Buffer B2 (1 M sodium perchlorate, 30% acetonitrile) at a flow rate of 10 ml/min for 30 min at 65°C. Purified oligonucleotides were collected, desalted by size-exclusion chromatography using a Sephadex G25 column (GE Healthcare Life Sciences, Marlborough, MA), and lyophilized.

### Analysis of oligonucleotides

Oligonucleotide purity and identity was determined by Liquid Chromatography-Mass Spectrometry (LC-MS) analysis on an Agilent 6530 accurate-mass Q-TOF LC/MS (Agilent technologies, Santa Clara, CA). Liquid chromatography was performed using a 2.1 × 50-mm AdvanceBio oligonucleotide column (Agilent technologies, Santa Clara, CA) and a gradient of buffers A (9 mM trimethylamine, 100 mM hexafluoroisopropanol in water) and B (9 mM triethylamine/100 mM hexafluoroisopropanol in methanol). Sense strand gradient: 1% B to 40% B from 0 to 2 min, 40% B to 100% B from 2 to 10.5 min. Antisense strand gradient: 1% B to 12% B from 0 to 2 min, 12% B to 30% B from 2 to 10.5 min, and 30% B to 100% B from 10.5 to 11 min. Relative hydrophobicities of conjugated siRNAs were determine by measuring the retention time on an Agilent Prostar System equipped with a Water HxSil C18 column (75 × 4.6) in a gradient of 100% buffer A (0.1 M trimethylamine acetate in water) to 100% buffer B (0.1 M trimethylamine acetate in acetonitrile) at a flow rate of 1 ml/min for 15 min at 60°C.

### Injection of conjugated siRNAs in mice

Animal experiments were performed in accordance with animal care ethics approval and guidelines of University of Massachusetts Medical School Institutional Animal Care and Use Committee (IACUC, protocol number A-2411). Female FB/NJ mice (The Jackson Laboratory) 7- to 8-weeks old were injected subcutaneously with phosphate buffered saline (PBS controls) or with 20 mg/kg siRNA (unconjugated or lipid-conjugated) suspended in PBS (160 µL). For distribution studies, 3 mice per conjugate were injected (n = 49, included controls). For the efficacy studies, 8 mice per conjugate, per gene were studied (n = 272, included controls). For the toxicity studies, 3 mice per conjugate were injected (n = 30, included controls).

### Fluorescence microscopy

At 48-hours post-injection, mice were euthanized and perfused with PBS. Tissues were collected and immersed in 10% formalin solution overnight at 4°C. Tissues were embedded in paraffin and sliced into 4-μm sections that were mounted on glass slides. Tissue sections on glass slides were deparaffinized by incubating twice in xylene for 8 min. Sections were rehydrated in an ethanol series from 100% to 95% to 80%, for 4 min each. Slides were then washed twice with PBS, 2 min each, incubated with DAPI (250 ng/mL, Molecular Probes) in PBS for 1 minute, and washed again in PBS for 2 minutes. Slides were mounted with PermaFluor mounting medium (Molecular Probes) coverslips, and dried overnight at 4°C. Sections were imaged at 5× and 40× using a Leica DM5500B microscope fitted with a DFC365 FX fluorescence camera.

### Peptide nucleic acid (PNA) hybridization assay

Tissue concentrations of antisense strands were determined using a PNA hybridization assay^37, 38^. Tissues (15 mg) were placed in QIAGEN Collection Microtubes holding 3-mm tungsten beads and lysed in 300 µl MasterPure tissue lysis solution (EpiCentre) containing 0.2 mg/ml proteinase K (Invitrogen) using a QIAGEN TissueLyser II. Lysates were then centrifuged at 1,000 × *g* for 10 min and incubated for 1 h at 55° to 60°C. Sodium dodecyl sulphate (SDS) was precipitated from lysates by adding 20 µL 3 M potassium chloride and pelleted centrifugation at 5000 × *g* for 15 minutes. Conjugated siRNAs in cleared supernatant were hybridized to a Cy3-labeled PNA probe fully complementary to the antisense strand (PNABio, Thousand Oaks, CA, USA). Samples were analyzed by HPLC (Agilent, Santa Clara, CA) over a DNAPac PA100 anion-exchange column (Thermo Fisher Scientific), in a gradient of sodium perchlorate, as follows: Buffer A: 50% water; 50% acetonitrile; 25 mM Tris-HCl, pH 8.5; 1 mM ethylenediaminetetraacetate. Buffer B: 800 mM sodium perchlorate in buffer A. Gradient conditions: 10% buffer B within 4 min, 50% buffer B for 1 min, and 50% to 100% buffer B within 5 minutes. Cy3 fluorescence was monitored and peaks integrated. Final concentrations were ascertained using calibration curves generated by spiking known quantities of lipid-conjugated siRNA into tissue lysates from an untreated animal. Spiked samples for calibration and experimental samples were processed and analyzed under the same laboratory conditions.

### In vivo mRNA silencing experiments

At 1-week post-injection, mice were euthanized and perfused with PBS. Tissues were collected and stored in RNAlater (Sigma) at 4°C overnight. mRNA was quantified using the QuantiGene 2.0 Assay (Affymetrix). 1.5-mm punches (3 punches per tissue) were placed in QIAGEN Collection Microtubes holding 3-mm tungsten beads and lysed in 300 μl Homogenizing Buffer (Affymetrix) containing 0.2 mg/ml proteinase K (Invitrogen) using a QIAGEN TissueLyser II. Samples were then centrifuged at 1,000 × *g* for 10 min and incubated for 1 h at 55° to 60°C. Lysates and diluted probe sets (mouse *Htt*, mouse *Ppib*, or mouse *Hprt*) were added to the bDNA capture plate and signal was amplified and detected as described by Coles *et al*^51^. Luminescence was detected on a Tecan M1000 (Tecan, Morrisville, NC, USA).

### Statistical analysis

Data were analyzed using GraphPad Prism 7.01 software (GraphPad Software, Inc., San Diego, CA). For each independent mouse experiment, the level of silencing was normalized to the mean of the control group (PBS group). Data were analyzed using non-parametric one-way ANOVA with Bonferroni’s test for multiple comparisons, with significance calculated relative to both PBS controls and non-targeting controls (*Ntc*).

## Acknowledgements

This work was supported by NIH grants [RO1GM10880304, and S10 OD020012]. We thank Khvorova lab members for insightful discussions and support, Alexander Sun for helping with the fluorescence microscopy images and Darryl Conte for helping with the manuscript writing and editing.

## Author Contributions

A.B. designed and A.B. and D.E. synthesized all compounds. A.B. and A.C conducted all *in vivo* experiments. R.H. performed PNA hybridization assay measurements. A.K. and A.B. came up with the concept and wrote the manuscript.

## References

1 Zhou, J., Shum, K.-T., Burnett, J. C. & Rossi, J. J. Nanoparticle-based delivery of RNAi therapeutics: progress and challenges. Pharmaceuticals 6, 85–107 (2013).

2 Nair, J. K. et al. Multivalent N-Acetylgalactosamine-Conjugated siRNA localizes in hepatocytes and elicits robust RNAi-mediated gene silencing. JACS 136, 16958–16961 (2014).

3 Zimmermann, T. S. et al. Clinical proof of concept for a novel hepatocyte-targeting GalNAc-siRNA conjugate. Mol. Ther. 25, 71–78 (2017).

4 Rajeev, K. G. et al. Hepatocyte-specific delivery of siRNAs conjugated to novel non-nucleosidic trivalent N-Acetylgalactosamine elicits robust gene silencing in vivo. ChemBioChem 16, 903–908 (2015).

5 Matsuda, S. et al. siRNA conjugates carrying sequentially assembled trivalent N-Acetylgalactosamine linked through nucleosides elicit robust gene silencing in vivo in hepatocytes. ACS Chem. Biol. 10, 1181–1187 (2015).

6 Prakash, T. P. et al. Targeted delivery of antisense oligonucleotides to hepatocytes using triantennary N-acetyl galactosamine improves potency 10-fold in mice. NAR 42, 8796–8807 (2014).

7 Tanowitz, M. et al. Asialoglycoprotein receptor 1 mediates productive uptake of N-acetylgalactosamine-conjugated and unconjugated phosphorothioate antisense oligonucleotides into liver hepatocytes. Nucleic Acids Res. 45, 12388–12400 (2017).

8 Huang, H. Preclinical and clinical advances of GalNAc-decorated nucleic acid therapeutics. Mol. Ther. Nucleic Acids 6, 116–132 (2017).

9 Akinc, A. et al. A combinatorial library of lipid-like materials for delivery of RNAi therapeutics. Nat. Biotechnol. 26, 561–569 (2008).

10 Dahlman, J. E. et al. In vivo endothelial siRNA delivery using polymeric nanoparticles with low molecular weight. Nat. Nanotechnol. 9, 648–655 (2014).

11 Wolfrum, C. et al. Mechanisms and optimization of in vivo delivery of lipophilic siRNAs. Nat. Biotechnol. 25, 1149–1157 (2007).

12 Soutschek, J. et al. Therapeutic silencing of an endogenous gene by systemic administration of modified siRNAs. Nature 432, 173–178 (2004).

13 Khan, T. et al. Silencing myostatin using cholesterol-conjugated siRNAs induces muscle growth. Mol. Ther. Nucleic Acids 5, e342 (2016).

14 DiFiglia, M. et al. Therapeutic silencing of mutant huntingtin with siRNA attenuates striatal and cortical neuropathology and behavioral deficits. PNAS 104, 17204–17209 (2007).

15 Deng, Y. et al. Transdermal delivery of siRNA through microneedle array. Sci. Rep. 6, 21422 (2016).

16 Wu, Y., et al. Cell host & microbe article durable protection from herpes simplex virus-2 transmission following intravaginal application of siRNAs targeting both a viral and host gene. Cell Host Microbe 5, 84–94 (2009).

17 Alterman, J. F. et al. Hydrophobically modified siRNAs silence Huntingtin mRNA in primary neurons and mouse brain. Mol. Ther. Nucleic Acids 4, e266 (2015).

18 Nishina, K. et al. Efficient in vivo delivery of siRNA to the Lliver by conjugation of α-tocopherol. Mol. Ther. 16, 734–740 (2008).

19 Hassler, M. R. et al. Comparison of partially and fully chemically-modified siRNA in conjugate-mediated delivery in vivo. Nucleic Acids Res. doi:10.1093/nar/gky037 (2018).

20 Nair, J. K. et al. Impact of enhanced metabolic stability on pharmacokinetics and pharmacodynamics of GalNAc–siRNA conjugates. Nucleic Acids Res. 45, 10969–10977 (2017).

21 Foster, D. J. et al. Advanced siRNA designs further improve in vivo performance of GalNAc-siRNA conjugates. Mol. Ther. 26, 708–717 (2018).

22 Allerson, C. R. et al. Fully 2‘-modified oligonucleotide duplexes with improved in vitro potency and stability compared to unmodified small interfering RNA. J. Med. Chem. 48, 901–904 (2005).

23 Nallagatla, S. R. & Bevilacqua, P. C. Nucleoside modifications modulate activation of the protein kinase PKR in an RNA structure-specific manner. RNA 14, 1201–1213 (2008).

24 Jackson, A. L. et al. Position-specific chemical modification of siRNAs reduces “off-target” transcript silencing. RNA 12, 1197–1205 (2006).

25 Geary, R. S., Norris, D., Yu, R. & Bennett, C. F. Pharmacokinetics, biodistribution and cell uptake of antisense oligonucleotides. Adv. Drug. Deliv. Rev. 87, 46–51 (2015).

26 Eckstein, F. Developments in RNA chemistry, a personal view. Biochimie 84, 841–848 (2002).

27 Ma, J. B. et al. Structural basis for 5’-end-specific recognition of guide RNA by the A. fulgidus Piwi protein. Nature 434, 666–670 (2005).

28 Frank, F., Sonenberg, N. & Nagar, B. Structural basis for 5’-nucleotide base-specific recognition of guide RNA by human AGO2. Nature 465, 818–822 (2010).

29 Haraszti, R. A. et al. 5΄-Vinylphosphonate improves tissue accumulation and efficacy of conjugated siRNAs in vivo. Nucleic Acids Res. 45, 7581–7592 (2017).

30 Parmar, R. et al. 5’-(E)-Vinylphosphonate: a stable phosphate mimic can improve the RNAi activity of siRNA–GalNAc conjugates. ChemBioChem 17, 987–989 (2016).

31 van Meer, G., Voelker, D. R. & Feigenson, G. W. Membrane lipids: where they are and how they behave. Nat. Rev. Mol. Cell Biol. 9, 112–124 (2008).

32 Phillips, G. B. & Dodges, J. T. Composition of phospholipids and of phospholipid fatty acids of human plasma *J*. Lipid Res. 8, 676–681 (1967).

33 Nikan, M. et al. Synthesis and evaluation of parenchymal retention and efficacy of a metabolically stable O-Phosphocholine-N-docosahexaenoyl-l-serine siRNA conjugate in mouse brain. Bioconjugate Chem. 28, 758–1766 (2017).

34 Morrissey, D. V. et al. Activity of stabilized short interfering RNA in a mouse model of hepatitis B virus replication. Hepatology 41, 1349–1356 (2005).

35 Harbort, J. et al. Sequence, chemical, and structural variation of small interfering RNAs and short hairpin RNAs and the effect on mammalian gene silencing. Antisense Nucl. Acid Drug Dev. 13, 83–105 (2003).

36 Smith, M. & Jungalwala, F. B. Reversed-phase high performance liquid chromatography of phosphatidylcholine: a simple method for determining relative hydrophobic interaction of various molecular species. J. Lipid Res. 22, 697–704 (1981).

37 Roehl, I., Schuster, M. & Seiffert, S. Oligonucleotide detection method. US Patent US20110201006A1, 1–9 (2011).

38 Godinho, B. M. D. C. et al. Pharmacokinetic profiling of conjugated therapeutic oligonucleotides: a high-throughput method based upon serial blood microsampling coupled to Peptide Nucleic Acid hybridization assay. Ncleic Acid Ther. 27, 323–334 (2017).

39 Oberbauer, R., Schreiner, G. F. & Meyer, T. W. Renal uptake of an 18-mer phosphorothioate oligonucleotide. Kidney Int. 48, 1226–1232 (1995).

40 Geary, R. S. Antisense oligonucleotide pharmacokinetics and metabolism. Expert Opin. Drug Metab. Toxicol. 5, 381–391 (2009).

41 Reed, D. R., Bachmanov, A. A. & Tordoff, M. G. Forty mouse strain survey of body composition. Physiol. Behav. 91, 593–600 (2007).

42 Wanke, R. et al. Overgrowth of skin in growth hormone transgenic mice depends on the presence of male gonads. J. Investig. Dermatol. 113, 967–971 (1999).

43 Taniguchi, T. et al. Plasmodium berghei ANKA causes intestinal malaria associated with dysbiosis. Sci. Rep. 5, 1–12 (2015).

44 Reynolds, A. et al. Rational siRNA design for RNA interference. Nat. Biotechnol. 22, 326–330 (2004).

45 Crooke, S. T., Wang, S., Vickers, T. A., Shen, W. & Liang, X.-H. Cellular uptake and trafficking of antisense oligonucleotides. Nat. Biotechnol. 35, 230–237 (2017).

46 Ly, S. et al. Visualization of self-delivering hydrophobically modified siRNA cellular internalization. Nucleic Acids Res. 45, 15–25 (2017).

47 Ansseau, E. et al. Antisense oligonucleotides used to target the DUX4 mRNA as therapeutic approaches in FaciosScapuloHumeral Muscular Dystrophy (FSHD). Genes 8, 1–21 (2017).

48 Suhr, O. B. et al. Efficacy and safety of patisiran for familial amyloidotic polyneuropathy: a phase II multi-dose study. Orphanet J. Rare Dis. 10, 1–9 (2015).

49 Chen, S. et al. Influence of particle size on the in vivo potency of lipid nanoparticle formulations of siRNA. J. Control. Release 235, 236–244 (2016).

50 Nikan, M. et al. Docosahexaenoic acid conjugation enhances distribution and safety of siRNA upon local administration in mouse brain. Mol. Ther. Nucleic Acids 5, e344 (2016).

51 Coles, A. H. et al. A high-throughput method for direct detection of therapeutic oligonucleotide-induced gene silencing in vivo. Nucleic Acid Ther. 26, 86–92 (2016).

