## Supplementary Materials for "Diverse lipid conjugates for functional extra-hepatic siRNA delivery *in vivo*"

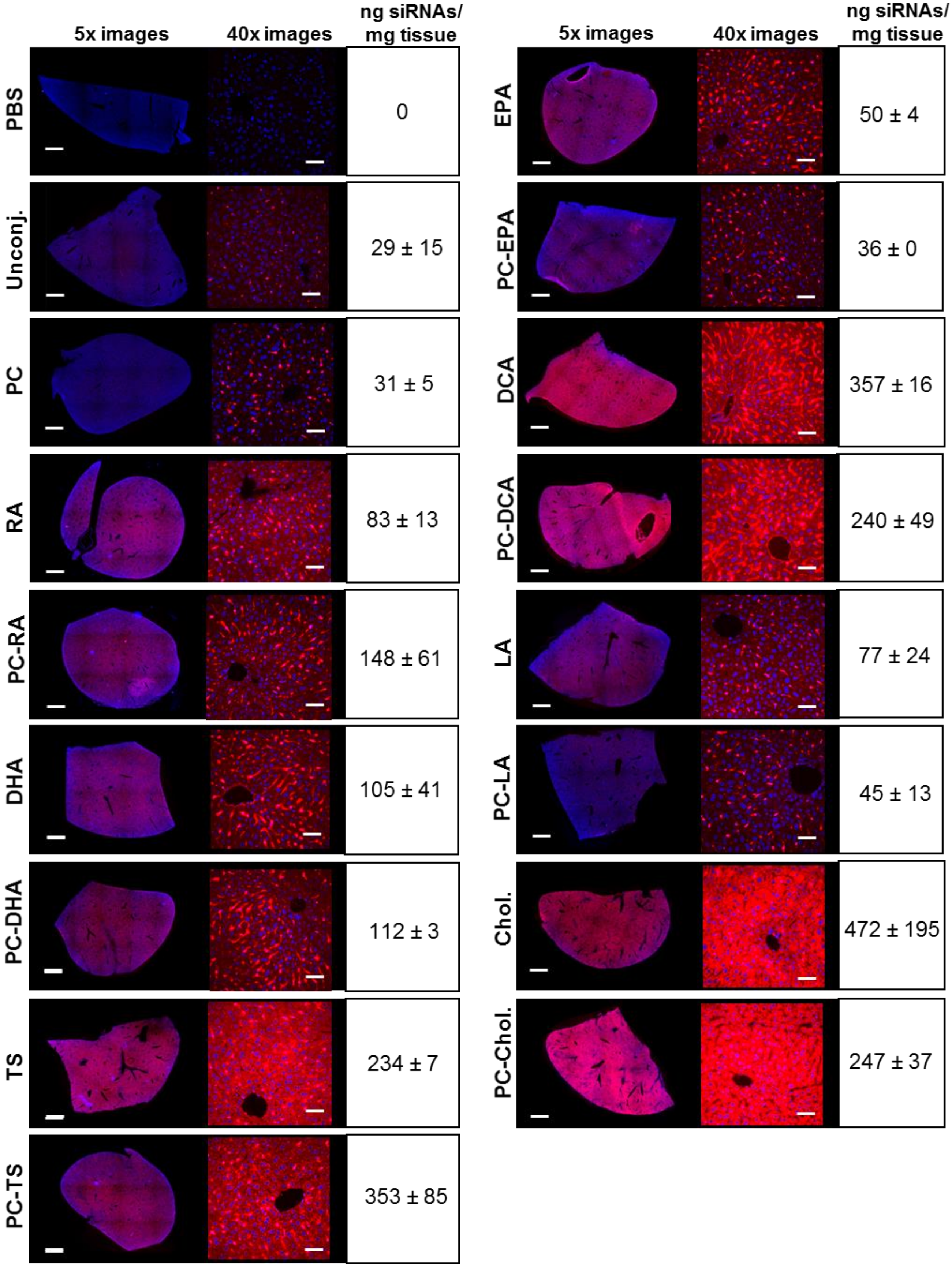


**Supplementary Figure 1: Liver distribution of Cy3-conjugated siRNAs.** Subcutaneous injection (FVB/N mice); 20 mg/kg; collection of tissues 48h after injection; n = 3 per conjugate. DAPI in blue; Cy3-siRNAs in red. 5x tiled arrays bar scale = 1 mm; 40x images bar scale = 50 µm; siRNA quantification by PNA hybridization assay (average of 3 animals ± SD).

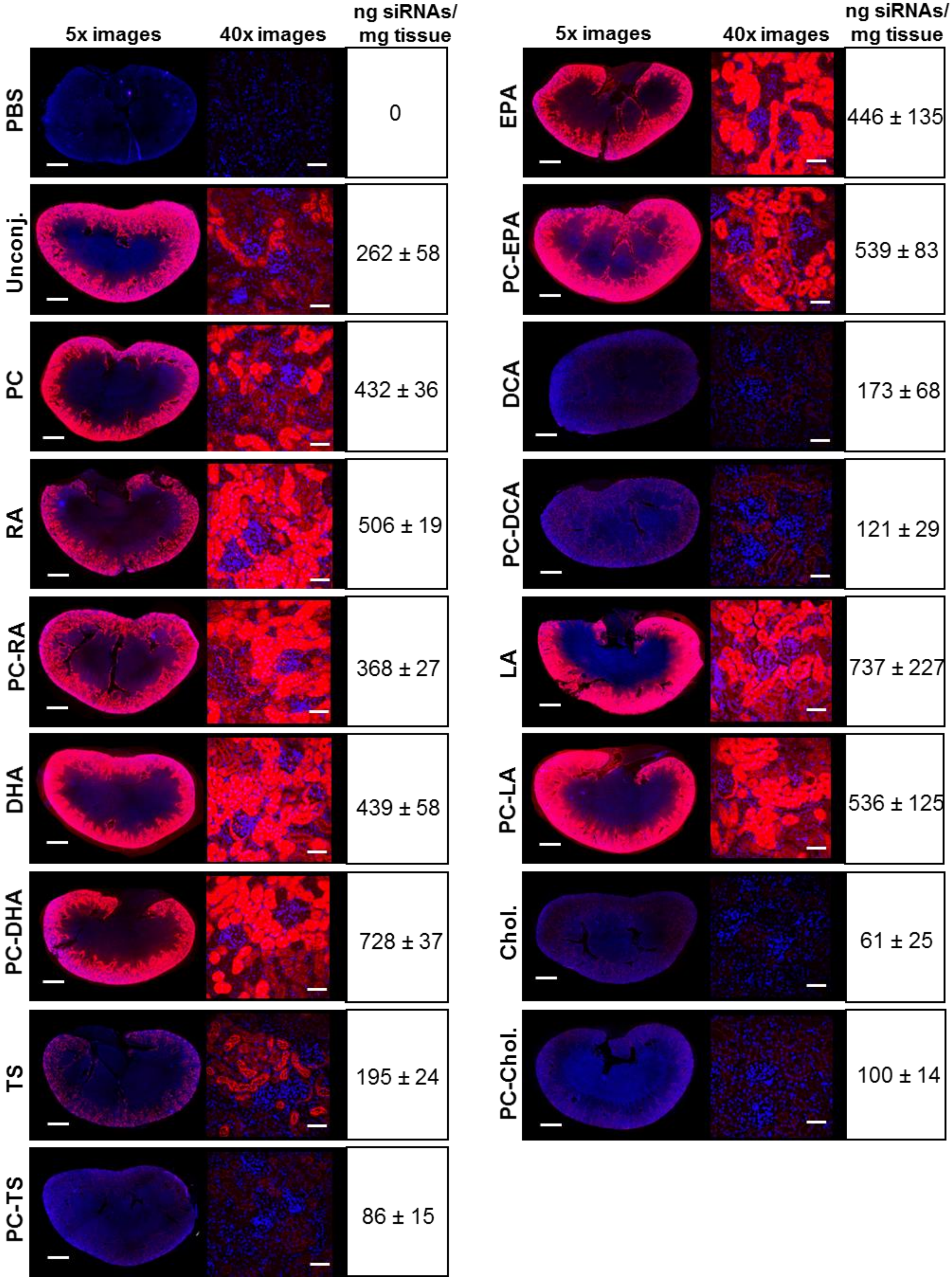


**Supplementary Figure 2: Kidney distribution of Cy3-conjugated siRNAs.** Subcutaneous injection (FVB/N mice); 20 mg/kg; collection of tissues 48h after injection; n = 3 per conjugate. DAPI in blue; Cy3-siRNAs in red. 5x tiled arrays bar scale = 1 mm; 40x images bar scale = 50 µm; siRNA quantification by PNA hybridization assay (average of 3 animals ± SD).

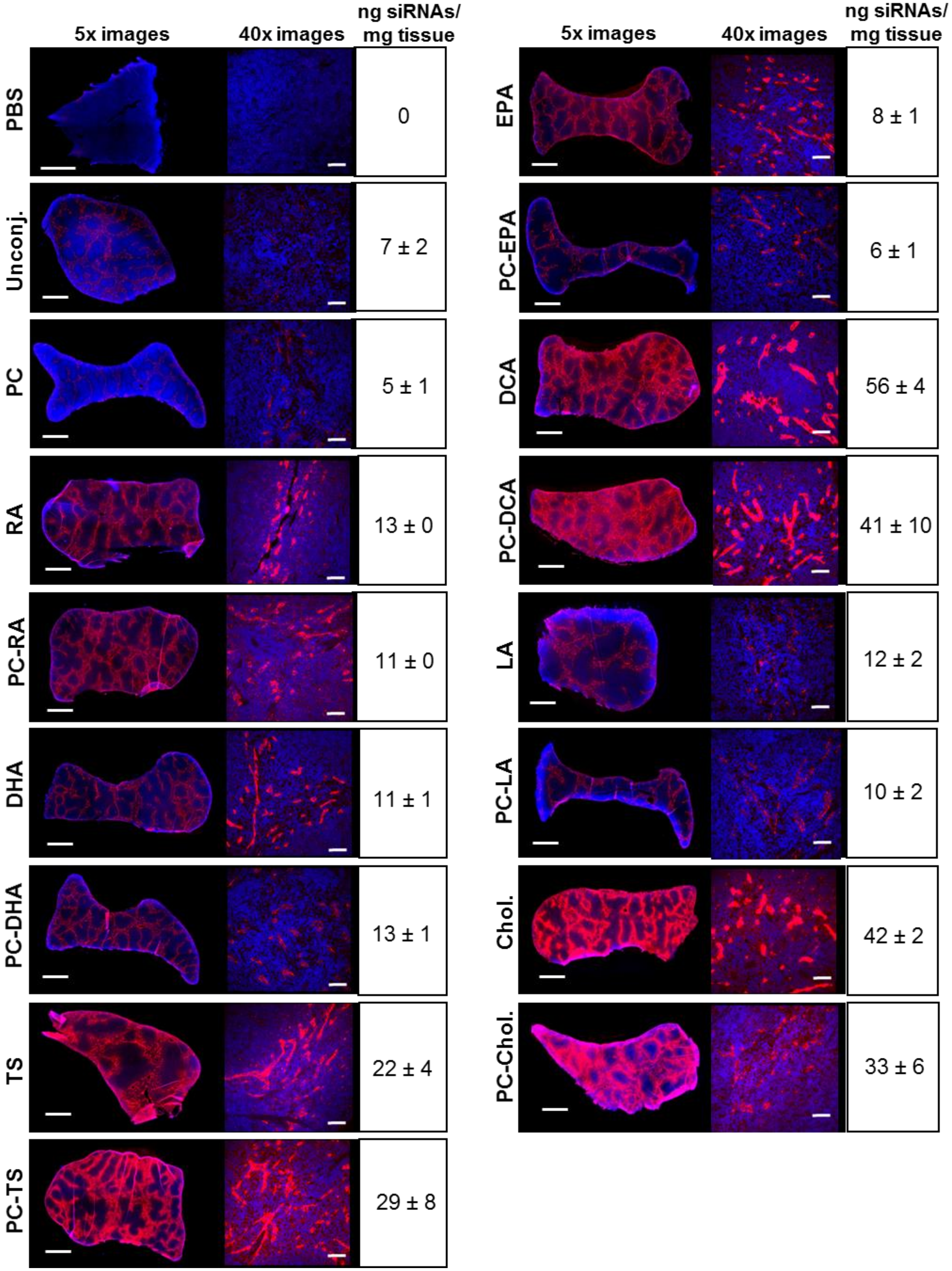


**Supplementary Figure 3: Spleen distribution of Cy3-conjugated siRNAs.** Subcutaneous injection (FVB/N mice); 20 mg/kg; collection of tissues 48h after injection; n = 3 per conjugate. DAPI in blue; Cy3-siRNAs in red. 5x tiled arrays bar scale = 1 mm; 40x images bar scale = 50 µm; siRNA quantification by PNA hybridization assay (average of 3 animals ± SD).

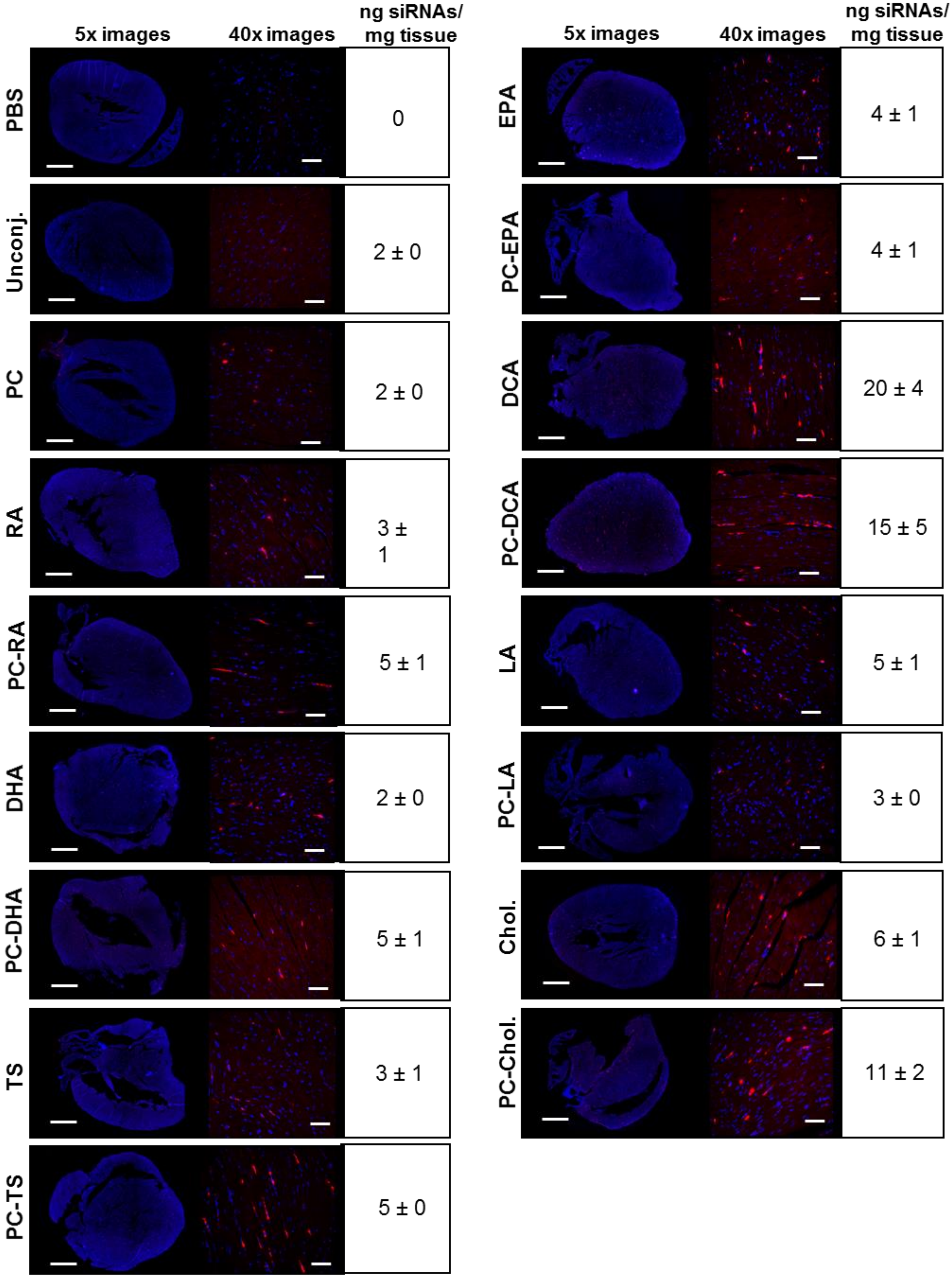


**Supplementary Figure 4: Heart distribution of Cy3-conjugated siRNAs.** Subcutaneous injection (FVB/N mice); 20 mg/kg; collection of tissues 48h after injection; n = 3 per conjugate. DAPI in blue; Cy3-siRNAs in red. 5x tiled arrays bar scale = 1 mm; 40x images bar scale = 50 µm; siRNA quantification by PNA hybridization assay (average of 3 animals ± SD).

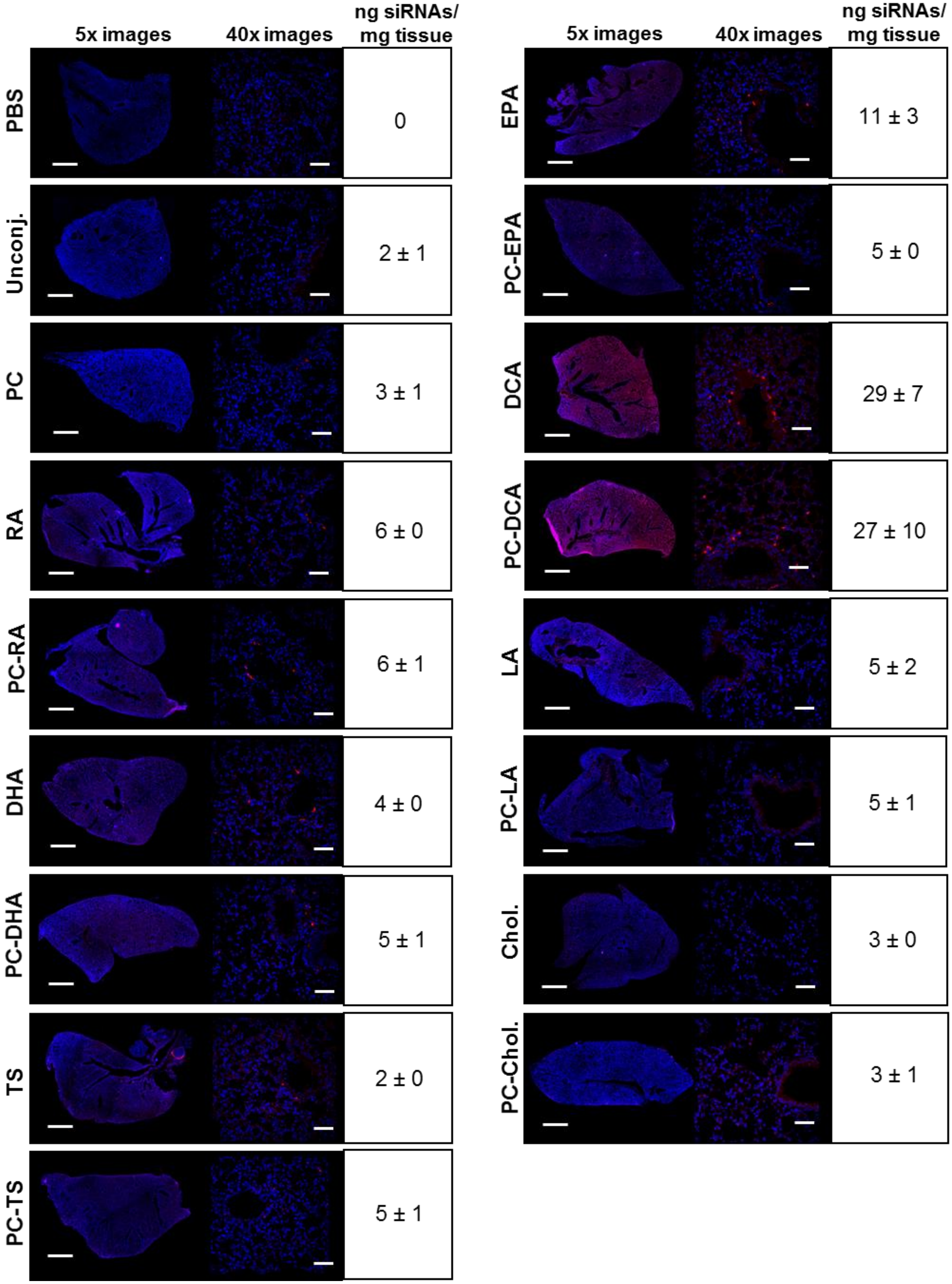


**Supplementary Figure 5: Lung distribution of Cy3-conjugated siRNAs.** Subcutaneous injection (FVB/N mice); 20 mg/kg; collection of tissues 48h after injection; n = 3 per conjugate. DAPI in blue; Cy3-siRNAs in red. 5x tiled arrays bar scale = 1 mm; 40x images bar scale = 50 µm; siRNA quantification by PNA hybridization assay (average of 3 animals ± SD).

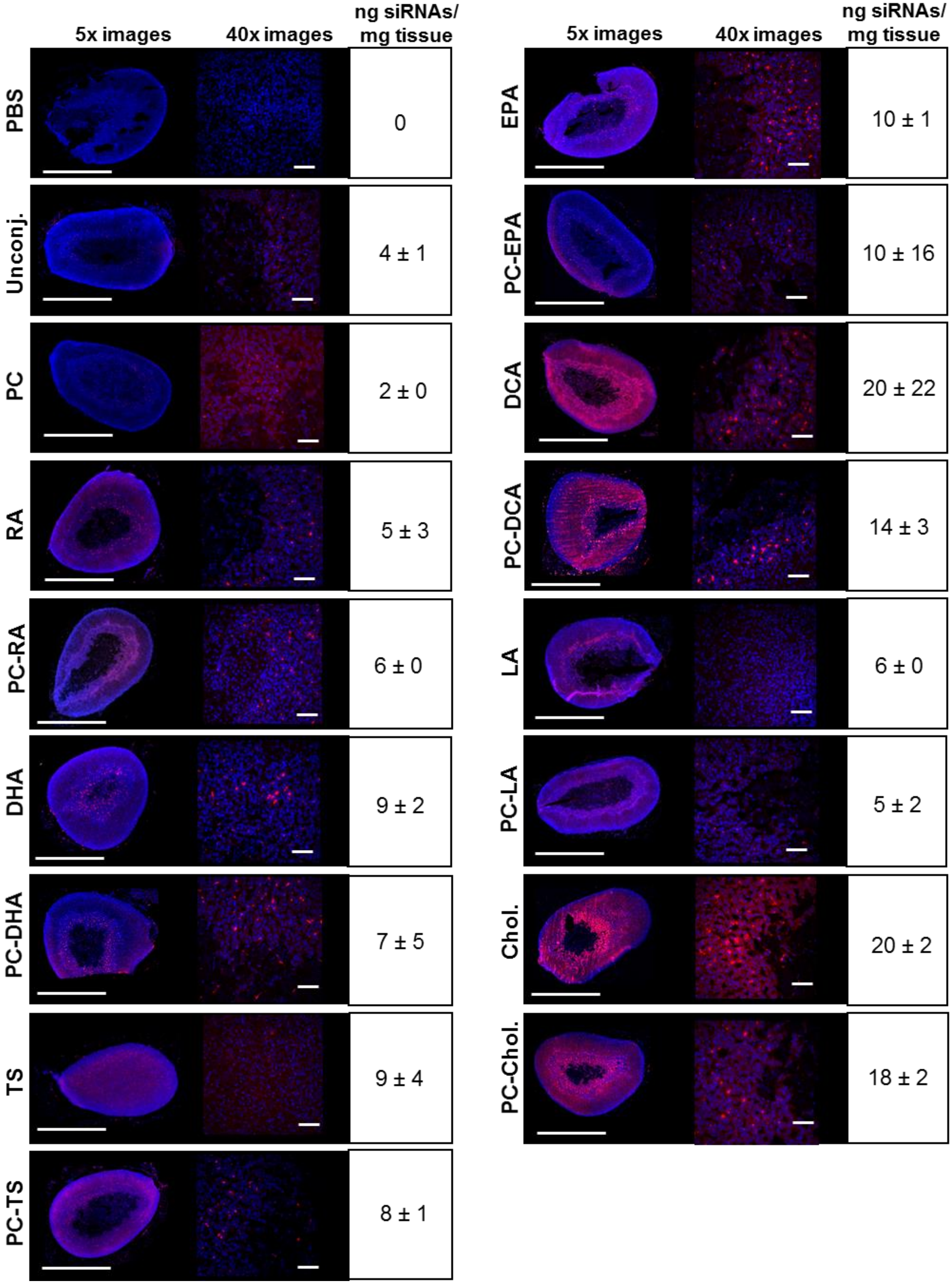


**Supplementary Figure 6: Adrenal glands distribution of Cy3-conjugated siRNAs.** Subcutaneous injection (FVB/N mice); 20 mg/kg; collection of tissues 48h after injection; n = 3 per conjugate. DAPI in blue; Cy3-siRNAs in red. 5x tiled arrays bar scale = 1 mm; 40x images bar scale = 50 µm; siRNA quantification by PNA hybridization assay (average of 3 animals ± SD).

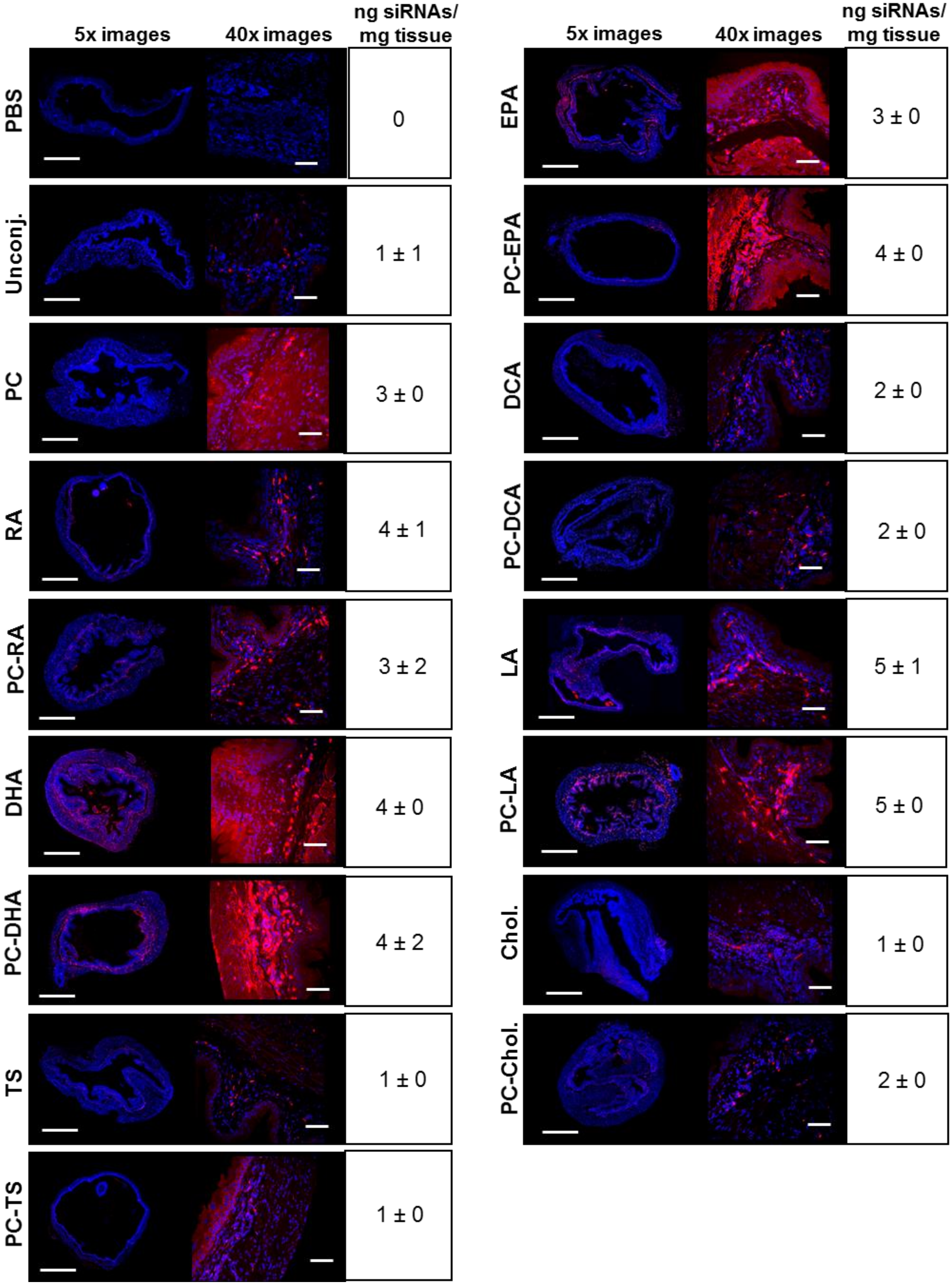


**Supplementary Figure 7: Bladder distribution of Cy3-conjugated siRNAs.** Subcutaneous injection (FVB/N mice); 20 mg/kg; collection of tissues 48h after injection; n = 3 per conjugate. DAPI in blue; Cy3-siRNAs in red. 5x tiled arrays bar scale = 1 mm; 40x images bar scale = 50 µm; siRNA quantification by PNA hybridization assay (average of 3 animals ± SD).

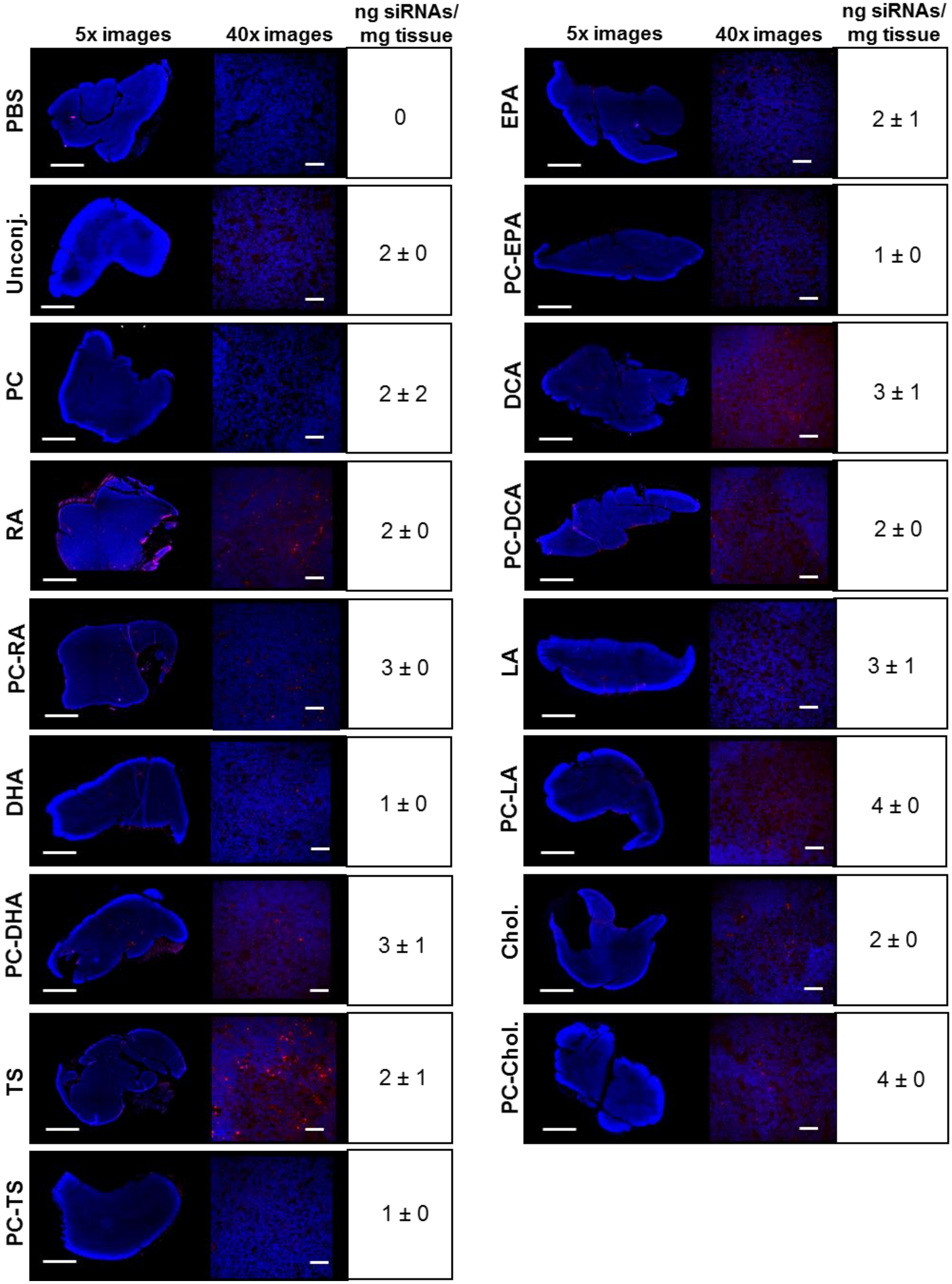


**Supplementary Figure 8: Thymus distribution of Cy3-conjugated siRNAs.** Subcutaneous injection (FVB/N mice); 20 mg/kg; collection of tissues 48h after injection; n = 3 per conjugate. DAPI in blue; Cy3-siRNAs in red. 5x tiled arrays bar scale = 1 mm; 40x images bar scale = 50 µm; siRNA quantification by PNA hybridization assay (average of 3 animals ± SD).

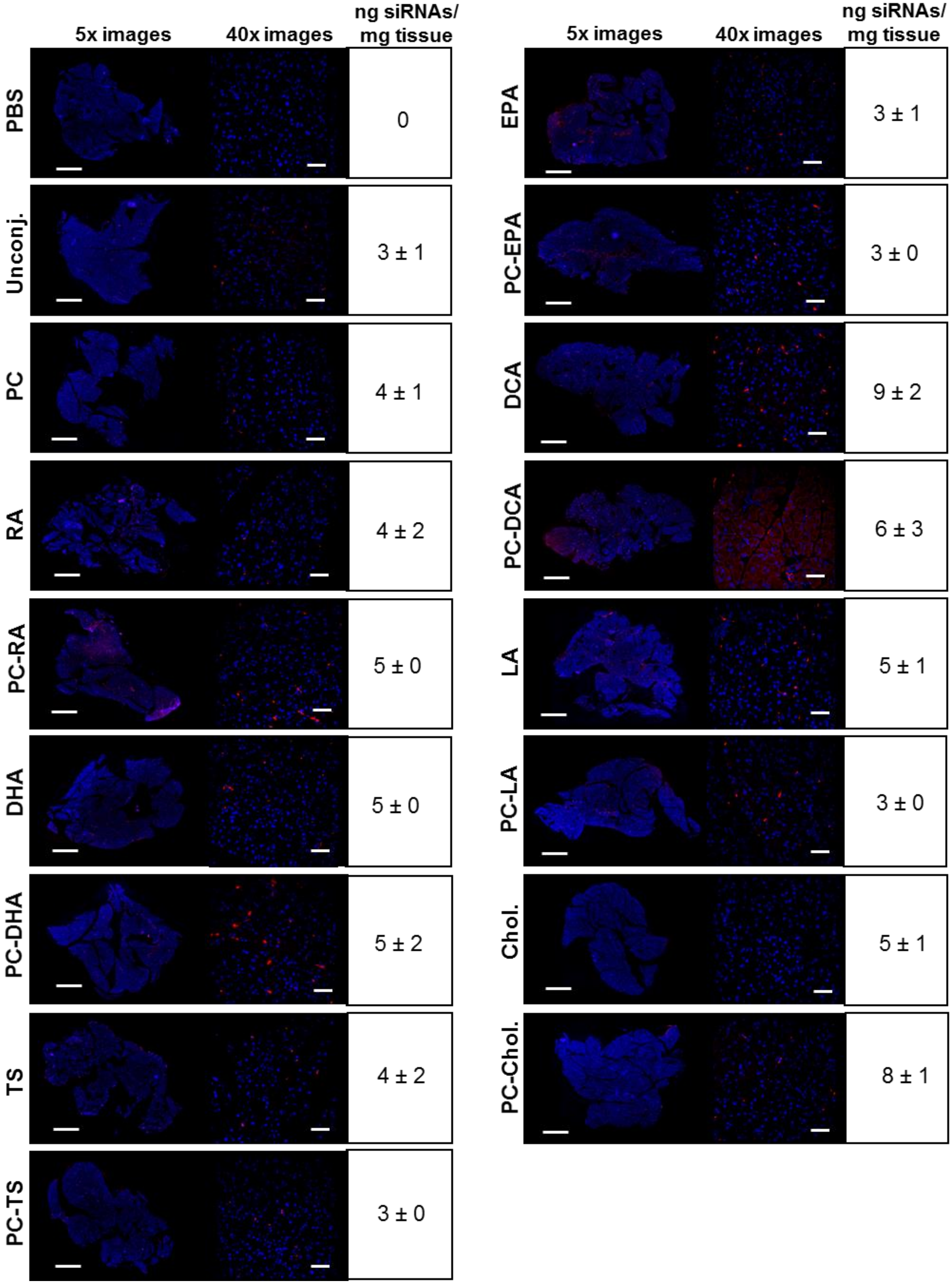


**Supplementary Figure 9: Pancreas distribution of Cy3-conjugated siRNAs.** Subcutaneous injection (FVB/N mice); 20 mg/kg; collection of tissues 48h after injection; n = 3 per conjugate. DAPI in blue; Cy3-siRNAs in red. 5x tiled arrays bar scale = 1 mm; 40x images bar scale = 50 µm; siRNA quantification by PNA hybridization assay (average of 3 animals ± SD).

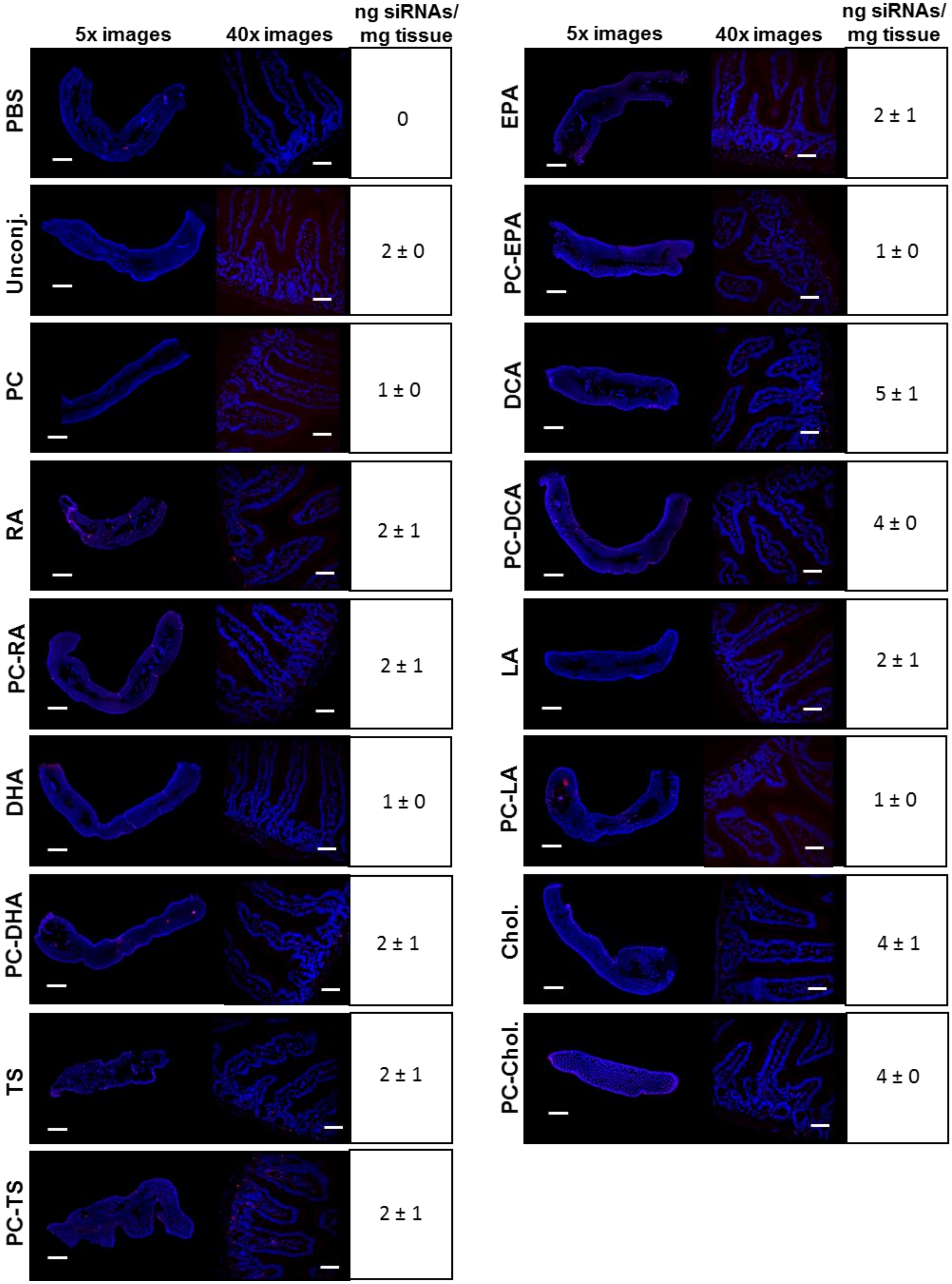


**Supplementary Figure 10: Intestine distribution of Cy3-conjugated siRNAs.** Subcutaneous injection (FVB/N mice); 20 mg/kg; collection of tissues 48h after injection; n = 3 per conjugate. DAPI in blue; Cy3-siRNAs in red. 5x tiled arrays bar scale = 1 mm; 40x images bar scale = 50 µm; siRNA quantification by PNA hybridization assay (average of 3 animals ± SD).

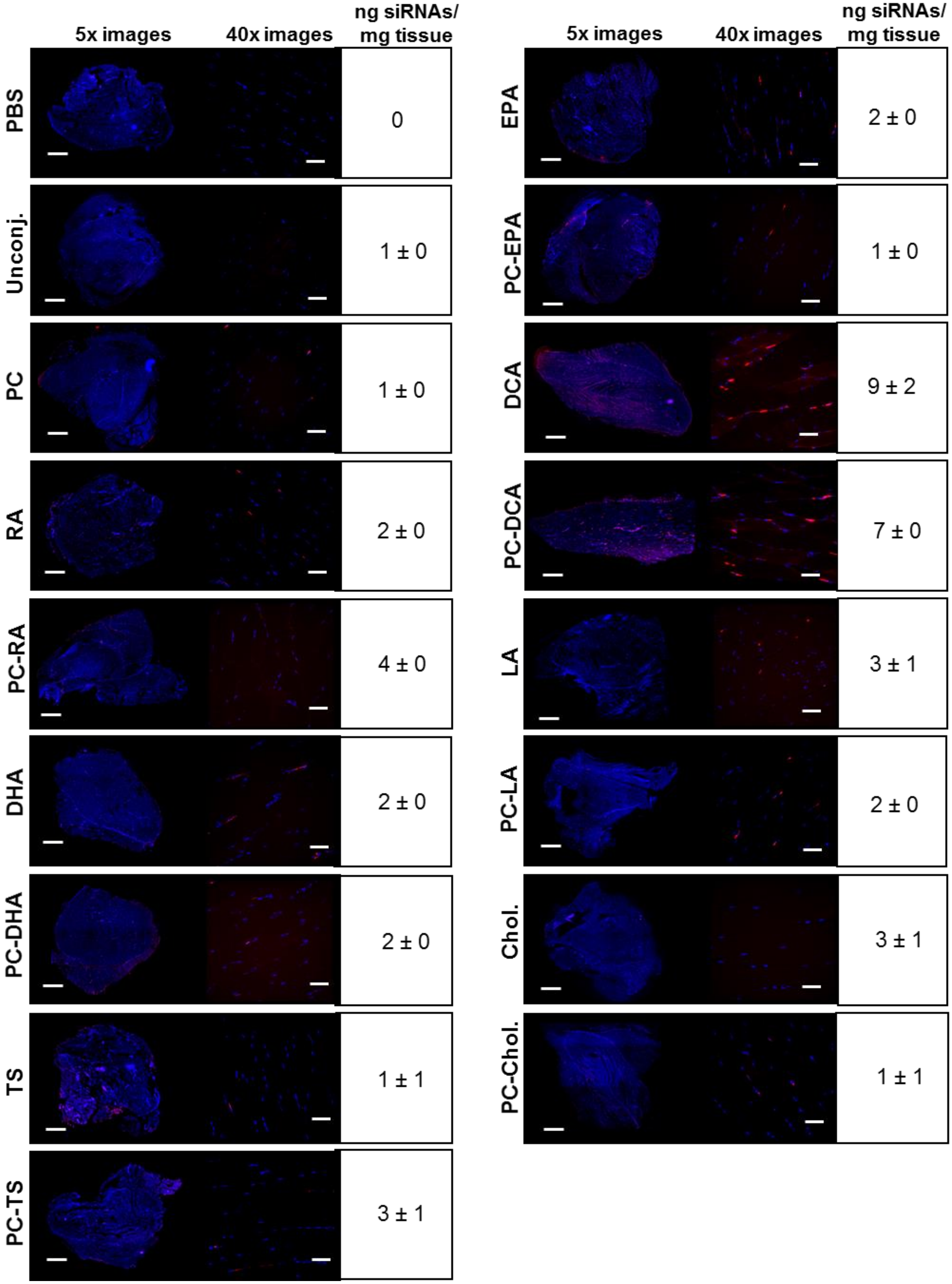


**Supplementary Figure 11: Muscle distribution of Cy3-conjugated siRNAs.** Subcutaneous injection (FVB/N mice); 20 mg/kg; collection of tissues 48h after injection; n = 3 per conjugate. DAPI in blue; Cy3-siRNAs in red. 5x tiled arrays bar scale = 1 mm; 40x images bar scale = 50 µm; siRNA quantification by PNA hybridization assay (average of 3 animals ± SD).

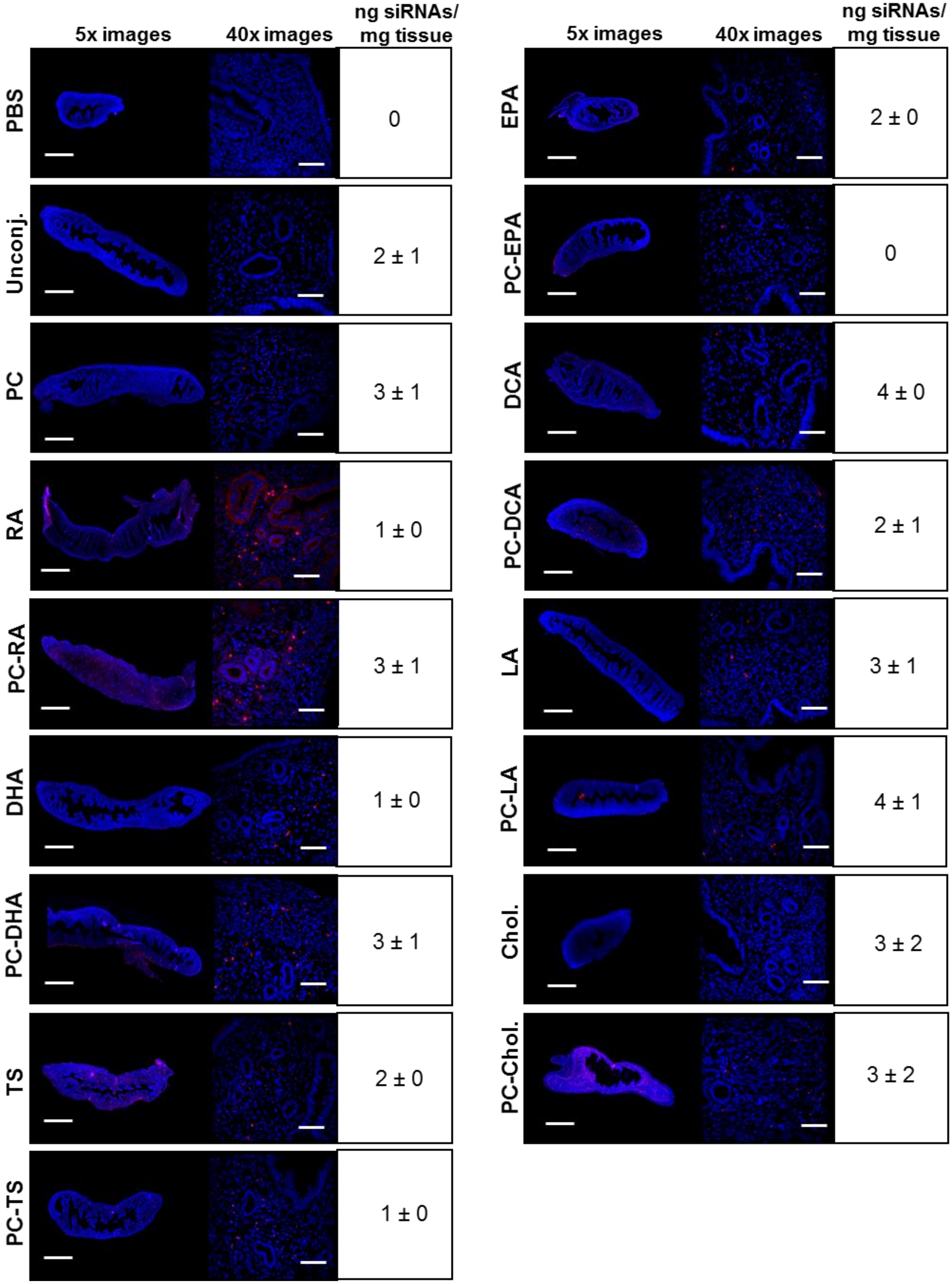


**Supplementary Figure 12: Fallopian tube distribution of Cy3-conjugated siRNAs.** Subcutaneous injection (FVB/N mice); 20 mg/kg; collection of tissues 48h after injection; n = 3 per conjugate. DAPI in blue; Cy3-siRNAs in red. 5x tiled arrays bar scale = 1 mm; 40x images bar scale = 50 µm; siRNA quantification by PNA hybridization assay (average of 3 animals ± SD).

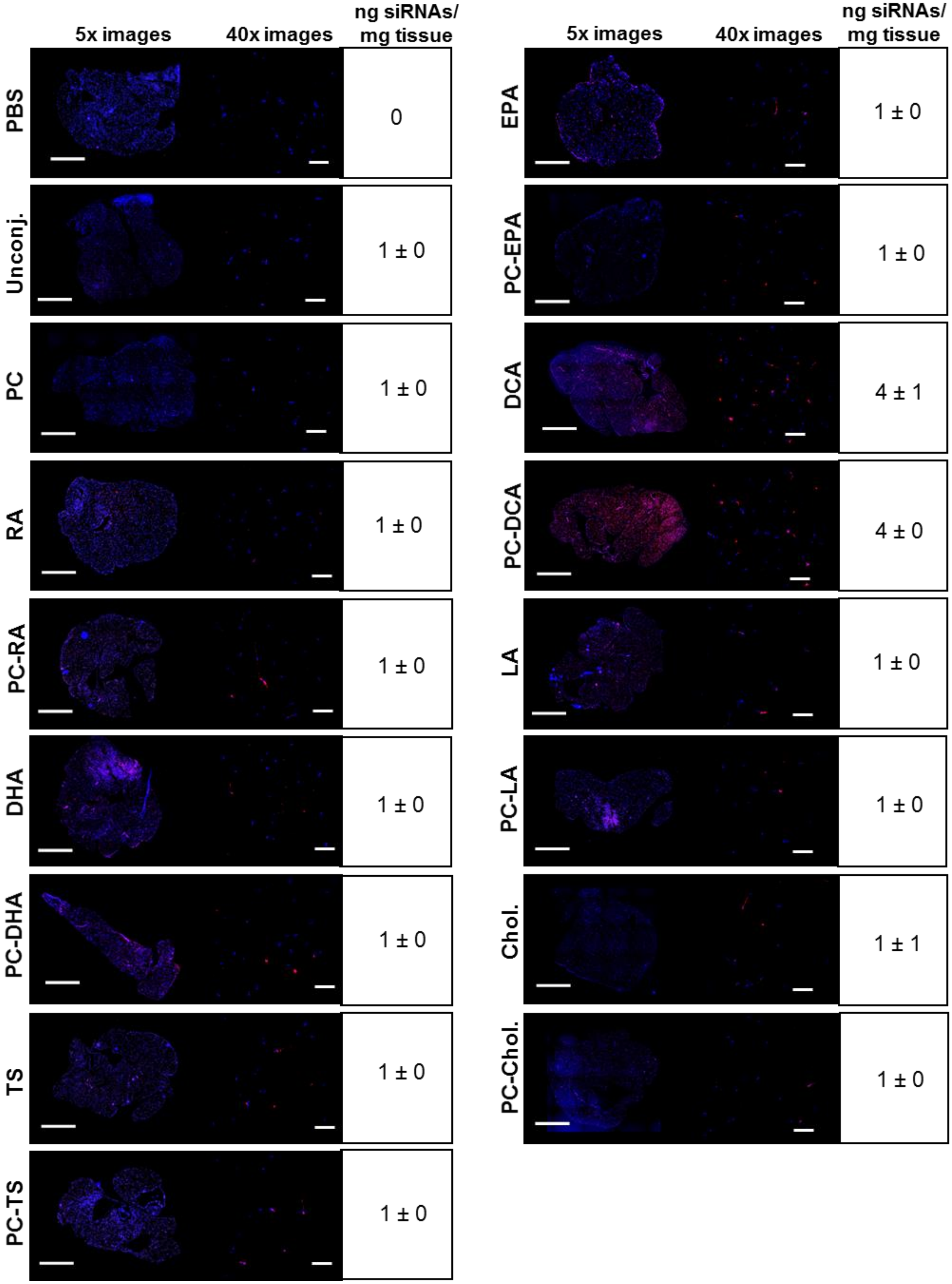


**Supplementary Figure 13: Fat distribution of Cy3-conjugated siRNAs.** Subcutaneous injection (FVB/N mice); 20 mg/kg; collection of tissues 48h after injection; n = 3 per conjugate. DAPI in blue; Cy3-siRNAs in red. 5x tiled arrays bar scale = 1 mm; 40x images bar scale = 50 μm; siRNA quantification by PNA hybridization assay (average of 3 animals ± SD).

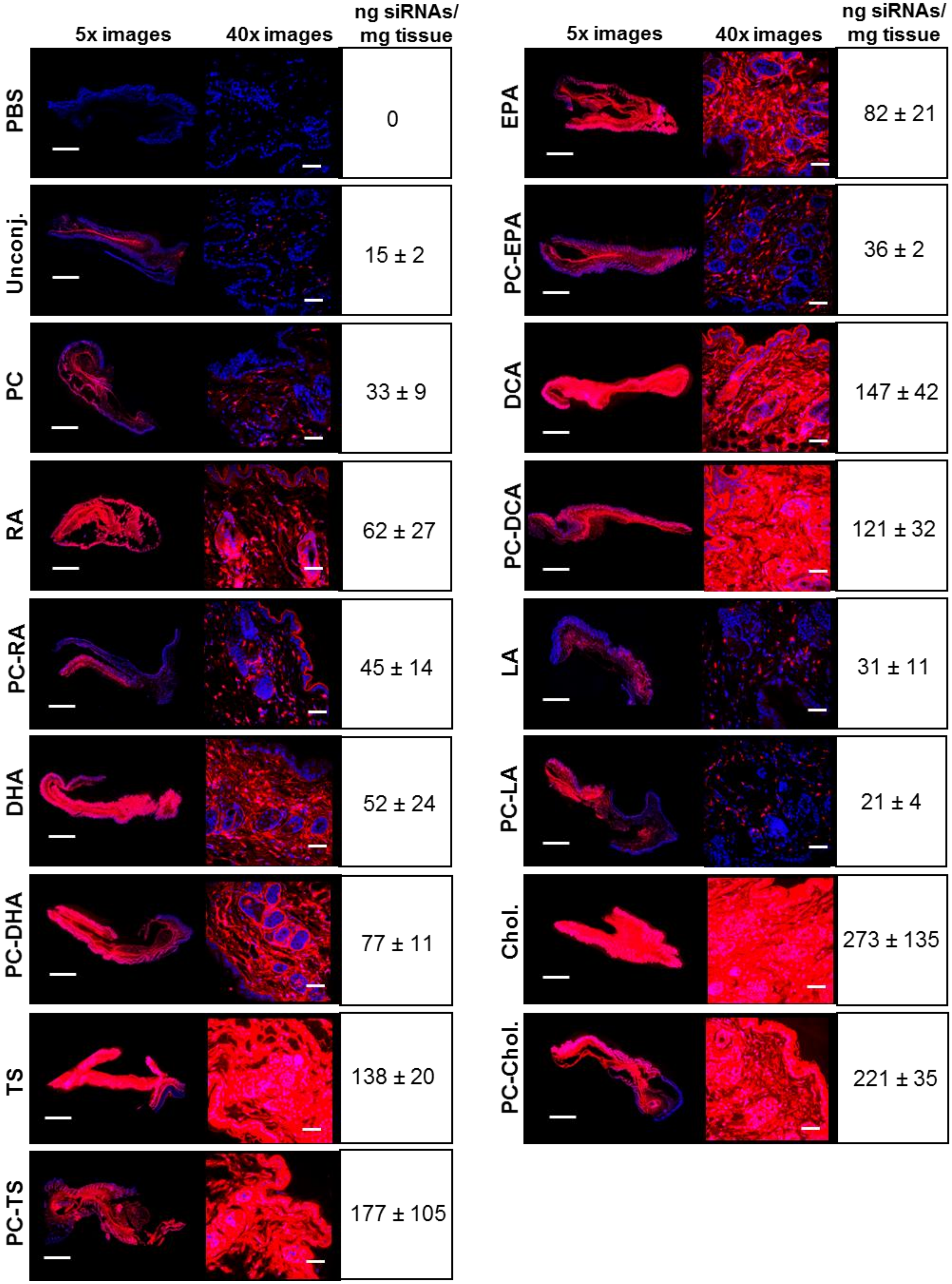


**Supplementary Figure 14: Skin (site of injection) of Cy3-conjugated siRNAs.** Subcutaneous injection (FVB/N mice); 20 mg/kg; collection of tissues 48h after injection; n = 3 per conjugate. DAPI in blue; Cy3-siRNAs in red. 5x tiled arrays bar scale = 1 mm; 40x images bar scale = 50 µm; siRNA quantification by PNA hybridization assay (average of 3 animals ± SD).

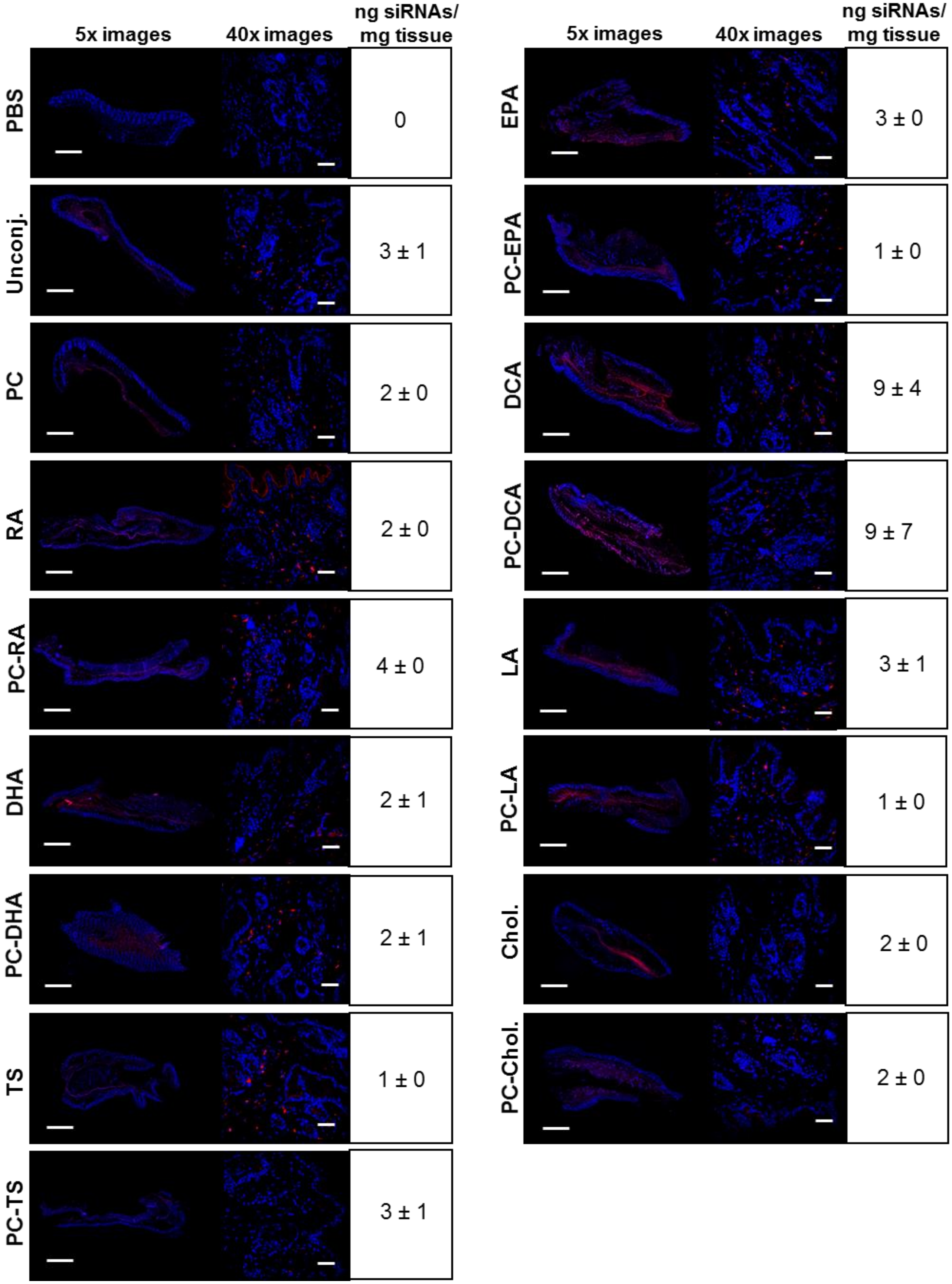


**Supplementary Figure 15: Skin of Cy3-conjugated siRNAs.** Subcutaneous injection (FVB/N mice); 20 mg/kg; collection of tissues 48h after injection; n = 3 per conjugate. DAPI in blue; Cy3-siRNAs in red. 5x tiled arrays bar scale = 1 mm; 40x images bar scale = 50 µm; siRNA quantification by PNA hybridization assay (average of 3 animals ± SD).

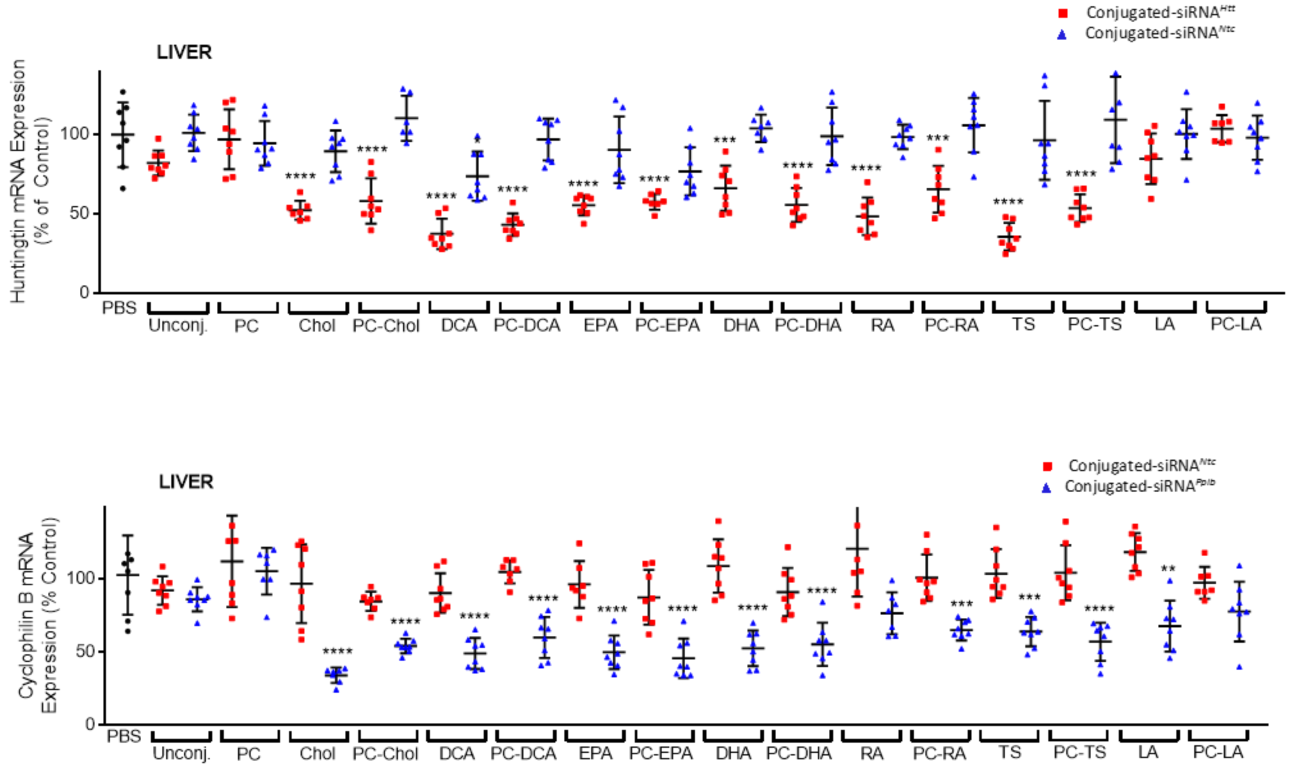


| | PBS | Unconj. | PC | Chol | PC-Chol | DCA | PC-DCA | EPA | PC-EPA | DHA | PC-DHA | RA | PC-RA | TS | PC-TS | LA | PC-LA |
|---|---|---|---|---|---|---|---|---|---|---|---|---|---|---|---|---|---|
| Huntingtin mRNA silencing (% compared to *Ntc*) | 0 ± 20 | 19 ± 8 | 0 ± 19 | 37 ± 6 | 52 ± 14 | 36 ± 9 | 54 ± 7 | 35 ± 6 | 19 ± 5 | 38 ± 14 | 43 ± 11 | 50 ± 12 | 40 ± 15 | 61 ± 9 | 56 ± 9 | 16 ± 16 | 0 ± 9 |
| Significance (compared to *Ntc*) | / | ns | ns | *** | **** | *** | **** | ** | ns | *** | **** | **** | **** | **** | **** | ns | ns |
| Cyclophilin B mRNA silencing (% compared to *Ntc*) | 0 ± 27 | 6 ± 8 | 7 ± 16 | 63 ± 5 | 30 ± 5 | 41 ± 11 | 45 ± 14 | 46 ± 12 | 41 ± 14 | 56 ± 12 | 36 ± 15 | 44 ± 14 | 36 ± 7 | 40 ± 10 | 47 ± 13 | 51 ± 17 | 20 ± 20 |
| Significance (compared to *Ntc*) | / | ns | ns | **** | ** | *** | *** | **** | *** | **** | ** | *** | ** | ** | **** | **** | ns |

**Supplementary Figure 16: Efficacy of conjugated siRNAs in liver.** Subcutaneous injection (FVB/N mice); 20 mg/kg; collection of tissues one week after injection; n = 16 per gene and per conjugate (included non-targeting controls or *Ntc*). Huntingtin (*Htt*) (upper panel) and Cyclophilin B (*Ppib*) (lower panel) mRNA levels were measured using QuantiGene® (Affymetrix), normalized to a housekeeping gene, *Hprt* (Hypoxanthine-guanine phosphoribosyl transferase), and presented as percent of PBS (Phosphate buffered saline) control (mean ± SD). Data analysis: Outliners define with Grubb's method (alpha = 0.1%); Multiple comparisons = One-way ANOVA, Bonferroni test (****P<0.0001, ***P<0.001, **P<0.01, *P<0.1). The table indicates the average of inhibition percentages (n = 8) and significances for each target and conjugate compared to *Ntc* (mean ± SD; ns = non-significant).

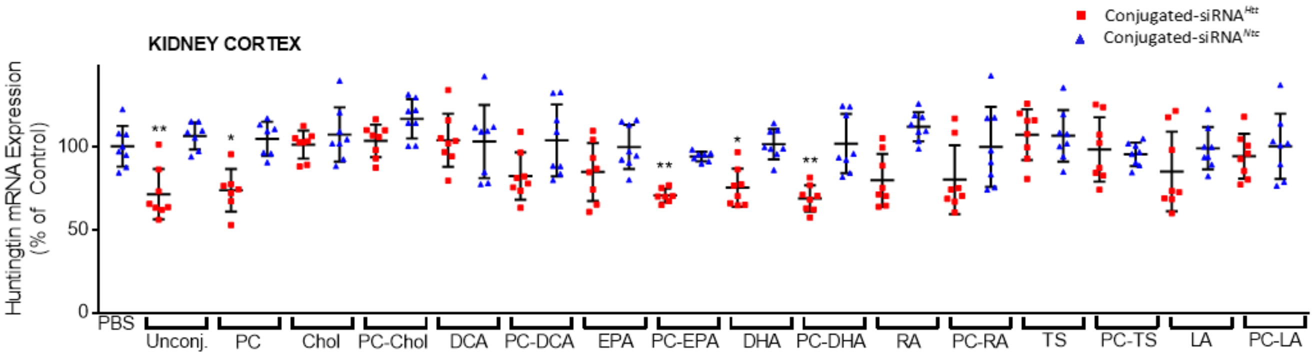


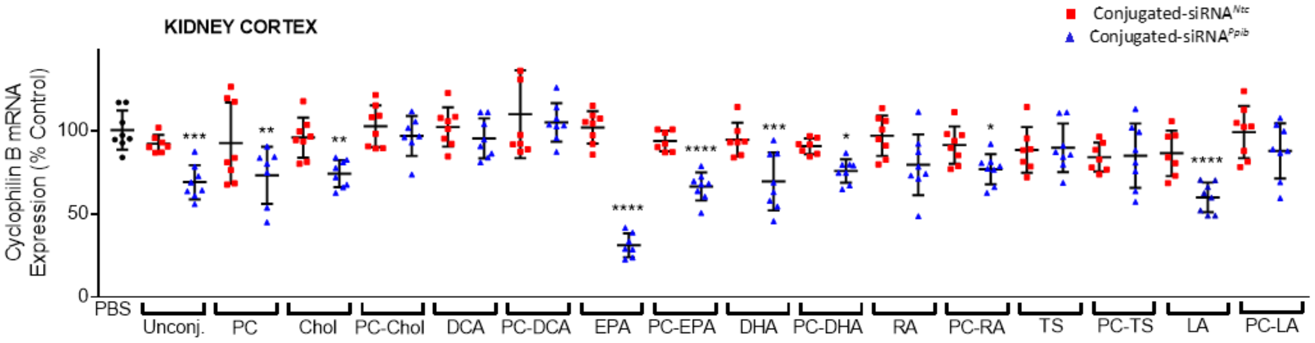


| | PBS | Unconj. | PC | Chol | PC-Chol | DCA | PC-DCA | EPA | PC-EPA | DHA | PC-DHA | RA | PC-RA | TS | PC-TS | LA | PC-LA |
|---|---|---|---|---|---|---|---|---|---|---|---|---|---|---|---|---|---|
| **Huntingtin mRNA silencing (% compared to *Ntc*)** | 0 ± 12 | 35 ± 15 | 31 ± 13 | 6 ± 8 | 13 ± 10 | 0 ± 16 | 22 ± 14 | 15 ± 17 | 24 ± 4 | 26 ± 11 | 33 ± 8 | 32 ± 16 | 20 ± 21 | 0 ± 15 | 0 ± 19 | 14 ± 24 | 6 ± 14 |
| **Significance (compared to *Ntc*)** | / | ** | * | ns | ns | ns | ns | ns | * | ns | ** | * | ns | ns | ns | ns | ns |
| **Cyclophilin B mRNA silencing (% compared to *Ntc*)** | 0 ± 12 | 23 ± 10 | 11 ± 24 | 22 ± 8 | 6 ± 12 | 7 ± 12 | 5 ± 12 | 71 ± 7 | 27 ± 8 | 25 ± 17 | 15 ± 7 | 18 ± 18 | 15 ± 9 | 0 ± 15 | 0 ± 19 | 27 ± 9 | 11 ± 17 |
| **Significance (compared to *Ntc*)** | / | * | ns | ns | ns | ns | ns | **** | ** | * | ns | ns | ns | ns | ns | * | ns |

**Supplementary Figure 17: Efficacy of conjugated siRNAs in kidney (cortex).** Subcutaneous injection (FVB/N mice); 20 mg/kg; collection of tissues one week after injection; n = 16 per gene and per conjugate (included non-targeting controls or *Ntc*). Huntingtin (*Htt*) (upper panel) and Cyclophilin B (*Ppib*) (lower panel) mRNA levels were measured using QuantiGene® (Affymetrix), normalized to a housekeeping gene, *Hprt* (Hypoxanthine-guanine phosphoribosyl transferase), and presented as percent of PBS (Phosphate buffered saline) control (mean ± SD). Data analysis: Outliners define with Grubb's method (alpha = 0.1%); Multiple comparisons = One-way ANOVA, Bonferroni test (****P<0.0001, ***P<0.001, **P<0.01, *P<0.1). The table indicates the average of inhibition percentages (n = 8) and significances for each target and conjugate compared to *Ntc* (mean ± SD; ns = non-significant).

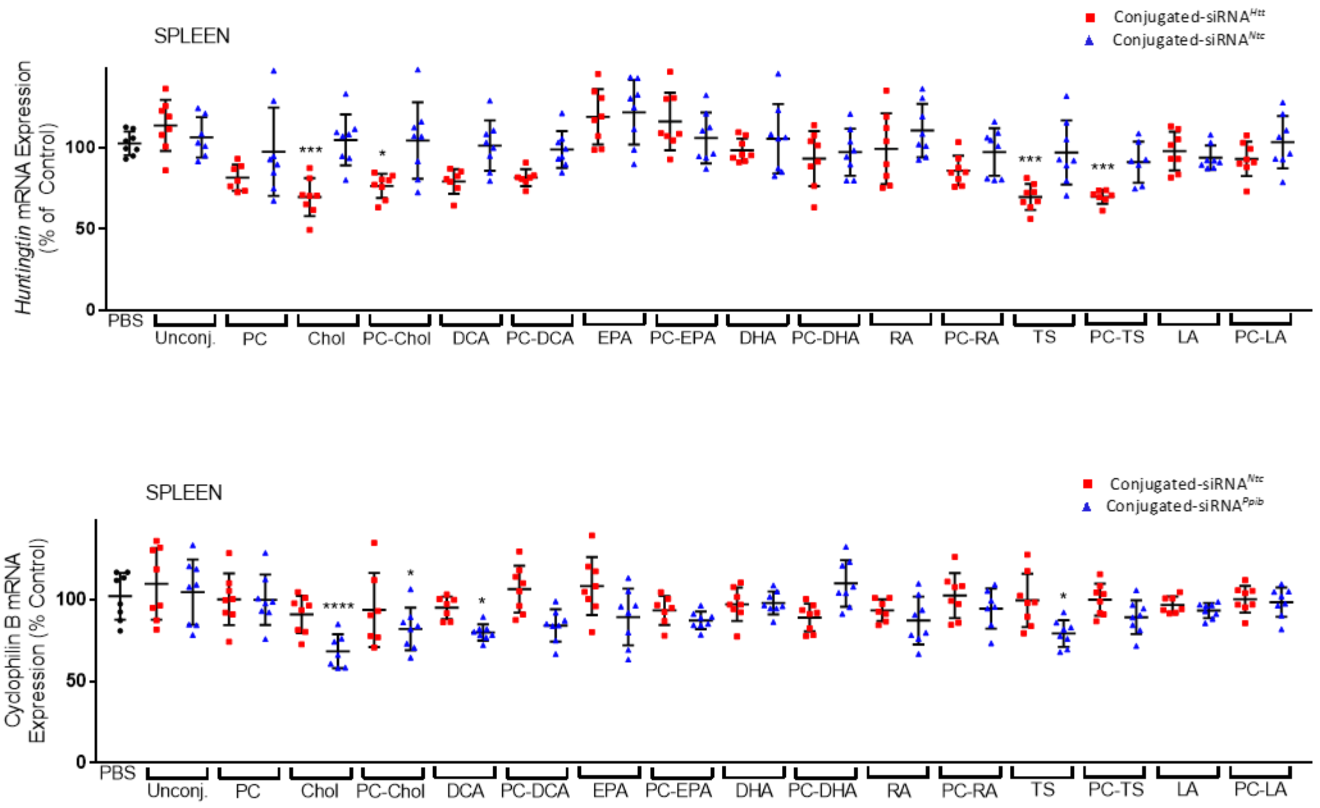


| | PBS | Unconj. | PC | Chol | PC-Chol | DCA | PC-DCA | EPA | PC-EPA | DHA | PC-DHA | RA | PC-RA | TS | PC-TS | LA | PC-LA |
|---|---|---|---|---|---|---|---|---|---|---|---|---|---|---|---|---|---|
| **Huntingtin mRNA silencing (% compared to *Ntc*)** | 0 ± 7 | 0 ± 16 | 16 ± 6 | 35 ± 12 | 28 ± 7 | 22 ± 8 | 17 ± 5 | 3 ± 17 | 0 ± 18 | 7 ± 7 | 4 ± 17 | 11 ± 22 | 11 ± 9 | 27 ± 8 | 21 ± 4 | 0 ± 12 | 10 ± 11 |
| **Significance (compared to *Ntc*)** | / | ns | ns | ** | ns | * | * | ns | ns | ns | ns | ns | ns | * | * | ns | ns |
| **Cyclophilin B mRNA silencing (% compared to *Ntc*)** | 0 ± 14 | 5 ± 20 | 0 ± 15 | 23 ± 10 | 12 ± 13 | 15 ± 5 | 22 ± 10 | 19 ± 17 | 6 ± 5 | 0 ± 7 | 0 ± 14 | 6 ± 15 | 8 ± 12 | 20 ± 8 | 11 ± 10 | 4 ± 5 | 2 ± 9 |
| **Significance (compared to *Ntc*)** | / | ns | ns | * | ns | * | ** | ns | ns | ns | ns | ns | ns | * | ns | ns | ns |

**Supplementary Figure 18: Efficacy of conjugated siRNAs in spleen.** Subcutaneous injection (FVB/N mice); 20 mg/kg; collection of tissues one week after injection; n = 16 per gene and per conjugate (included non-targeting controls or *Ntc*). Huntingtin (*Htt*) (upper panel) and Cyclophilin B (*Ppib*) (lower panel) mRNA levels were measured using QuantiGene® (Affymetrix), normalized to a housekeeping gene, *Hprt* (Hypoxanthine-guanine phosphoribosyl transferase), and presented as percent of PBS (Phosphate buffered saline) control (mean ± SD). Data analysis: Outliners define with Grubb's method (alpha = 0.1%); Multiple comparisons = One-way ANOVA, Bonferroni test (****P<0.0001, ***P<0.001, **P<0.01, *P<0.1). The table indicates the average of inhibition percentages (n = 8) and significances for each target and conjugate compared to *Ntc* (mean ± SD; ns = non-significant).

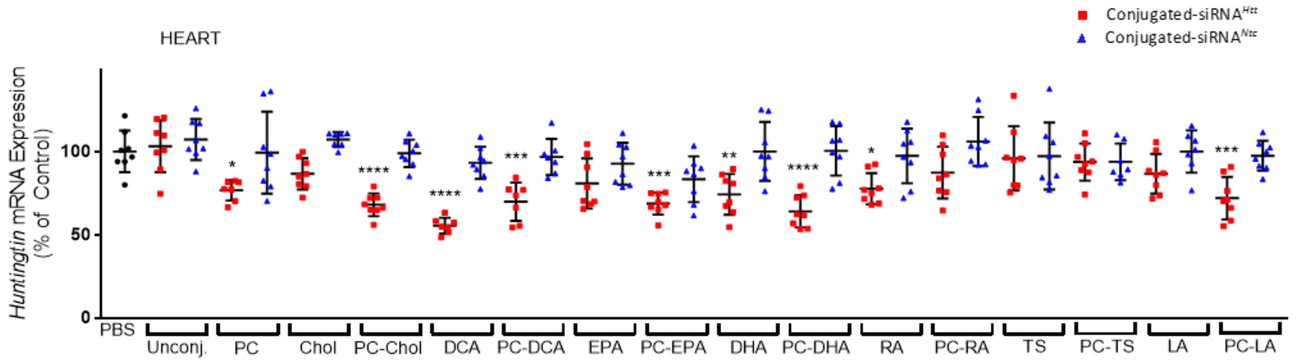


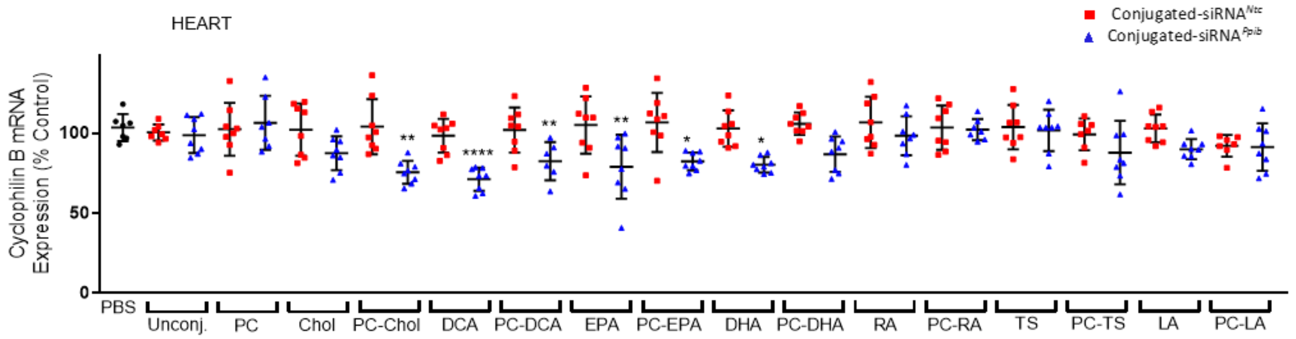


| | PBS | Unconj. | PC | Chol | PC-Chol | DCA | PC-DCA | EPA | PC-EPA | DHA | PC-DHA | RA | PC-RA | TS | PC-TS | LA | PC-LA |
|---|---|---|---|---|---|---|---|---|---|---|---|---|---|---|---|---|---|
| **Huntingtin mRNA silencing (% compared to *Ntc*)** | 0 ± 12 | 4 ± 16 | 23 ± 6 | 21 ± 9 | 31 ± 7 | 38 ± 5 | 20 ± 13 | 12 ± 15 | 15 ± 7 | 26 ± 12 | 36 ± 10 | 20 ± 9 | 19 ± 16 | 1 ± 19 | 0 ± 11 | 13 ± 12 | 25 ± 13 |
| **Significance (compared to *Ntc*)** | / | ns | ns | * | ** | **** | ** | ns | ns | ** | **** | ns | ns | ns | ns | ns | * |
| **Cyclophilin B mRNA silencing (% compared to *Ntc*)** | 0 ± 8 | 2 ± 11 | 0 ± 17 | 15 ± 11 | 29 ± 7 | 27 ± 7 | 11 ± 19 | 26 ± 20 | 24 ± 5 | 23 ± 5 | 19 ± 11 | 8 ± 12 | 1 ± 7 | 2 ± 13 | 11 ± 20 | 13 ± 6 | 1 ± 15 |
| **Significance (compared to *Ntc*)** | / | ns | ns | ns | ** | ** | ns | * | ns | ns | ns | ns | ns | ns | ns | ns | ns |

**Supplementary Figure 19: Efficacy of conjugated siRNAs in heart.** Subcutaneous injection (FVB/N mice); 20 mg/kg; collection of tissues one week after injection; n = 16 per gene and per conjugate (included non-targeting controls or *Ntc*). Huntingtin (*Htt*) (upper panel) and Cyclophilin B (*Ppib*) (lower panel) mRNA levels were measured using QuantiGene® (Affymetrix), normalized to a housekeeping gene, *Hprt* (Hypoxanthine-guanine phosphoribosyl transferase), and presented as percent of PBS (Phosphate buffered saline) control (mean ± SD). Data analysis: Outliners define with Grubb's method (alpha = 0.1%); Multiple comparisons = One-way ANOVA, Bonferroni test (****P<0.0001, ***P<0.001, **P<0.01, *P<0.1). The table indicates the average of inhibition percentages (n = 8) and significances for each target and conjugate compared to *Ntc* (mean ± SD; ns = non-significant).

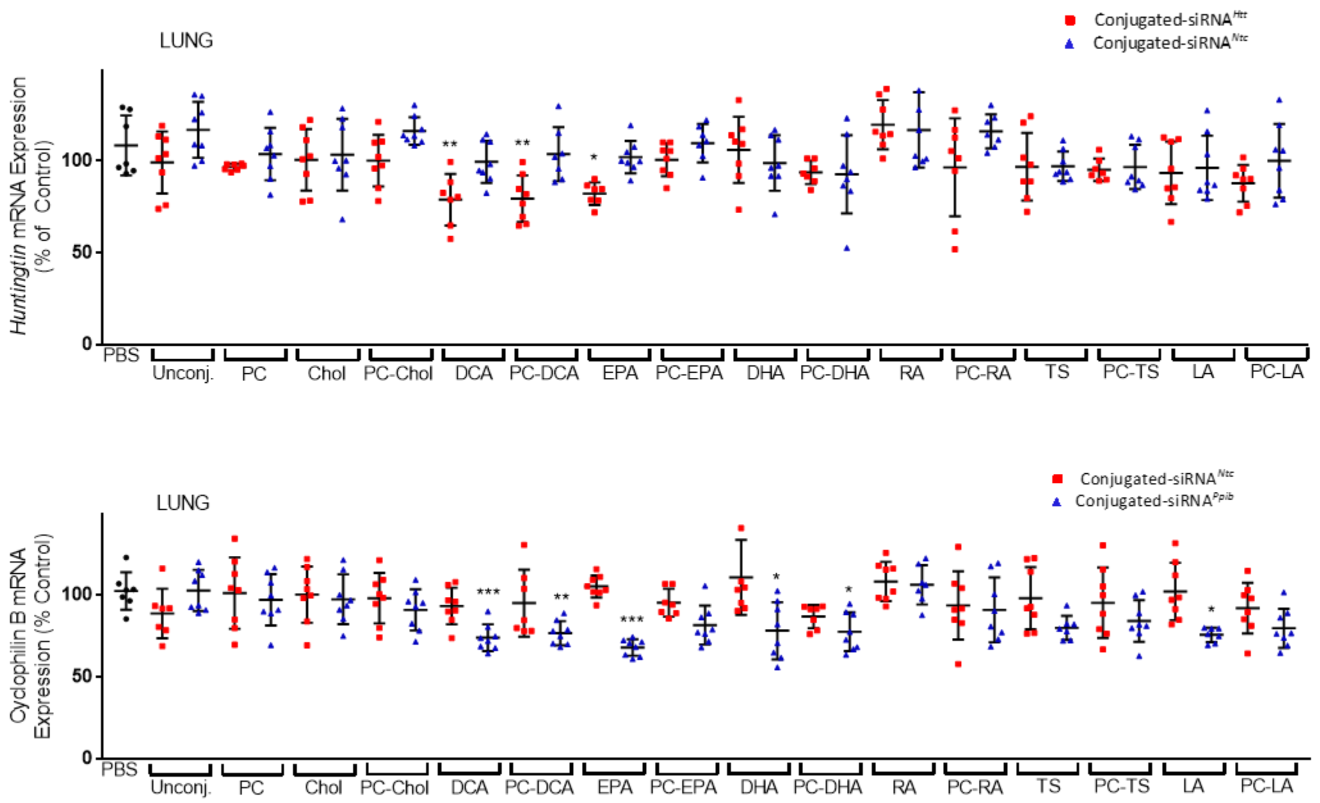


| | PBS | Unconj. | PC | Chol | PC-Chol | DCA | PC-DCA | EPA | PC-EPA | DHA | PC-DHA | RA | PC-RA | TS | PC-TS | LA | PC-LA |
|---|---|---|---|---|---|---|---|---|---|---|---|---|---|---|---|---|---|
| **Huntingtin mRNA silencing (% compared to *Ntc*)** | 0 ± 16 | 18 ± 17 | 7 ± 2 | 3 ± 17 | 16 ± 14 | 18 ± 15 | 24 ± 13 | 20 ± 6 | 9 ± 9 | 0 ± 18 | 0 ± 6 | 0 ± 13 | 20 ± 27 | 0 ± 18 | 2 ± 6 | 3 ± 17 | 12 ± 10 |
| **Significance (compared to *Ntc*)** | / | ns | ns | ns | ns | * | * | * | ns | ns | ns | ns | ns | ns | ns | ns | ns |
| **Cyclophilin B mRNA silencing (% compared to *Ntc*)** | 0 ± 11 | 0 ± 13 | 4 ± 16 | 3 ± 15 | 7 ± 13 | 19 ± 8 | 15 ± 7 | 35 ± 5 | 14 ± 12 | 33 ± 17 | 9 ± 12 | 2 ± 12 | 3 ± 20 | 18 ± 7 | 11 ± 12 | 27 ± 4 | 12 ± 12 |
| **Significance (compared to *Ntc*)** | / | ns | ns | ns | ns | * | ns | *** | ns | ** | ns | ns | ns | ns | ns | ns | ns |

**Supplementary Figure 20: Efficacy of conjugated siRNAs in lung.** Subcutaneous injection (FVB/N mice); 20 mg/kg; collection of tissues one week after injection; n = 16 per gene and per conjugate (included non-targeting controls or *Ntc*). Huntingtin (*Htt*) (upper panel) and Cyclophilin B (*Ppib*) (lower panel) mRNA levels were measured using QuantiGene® (Affymetrix), normalized to a housekeeping gene, *Hprt* (Hypoxanthine-guanine phosphoribosyl transferase), and presented as percent of PBS (Phosphate buffered saline) control (mean ± SD). Data analysis: Outliners define with Grubb's method (alpha = 0.1%); Multiple comparisons = One-way ANOVA, Bonferroni test (***P<0.001, **P<0.01, *P<0.1). The table indicates the average of inhibition percentages (n = 8) and significances for each target and conjugate compared to *Ntc* (mean ± SD; ns = non-significant).

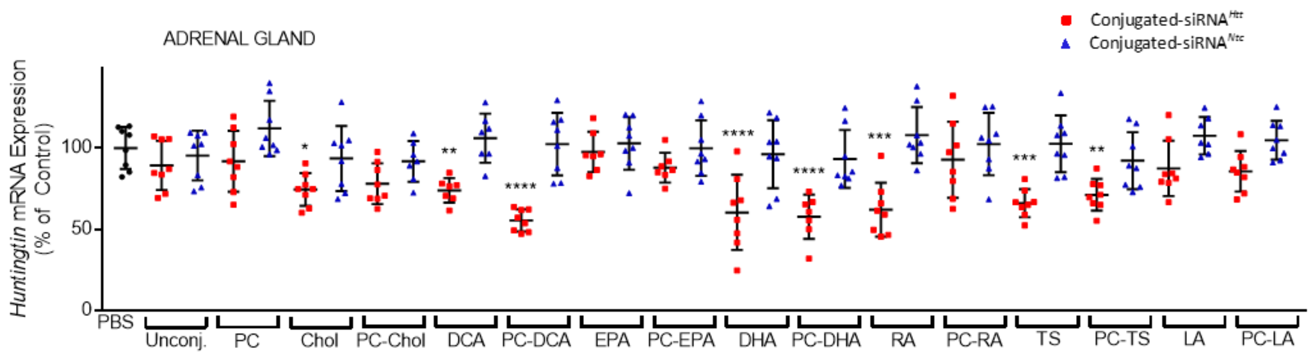


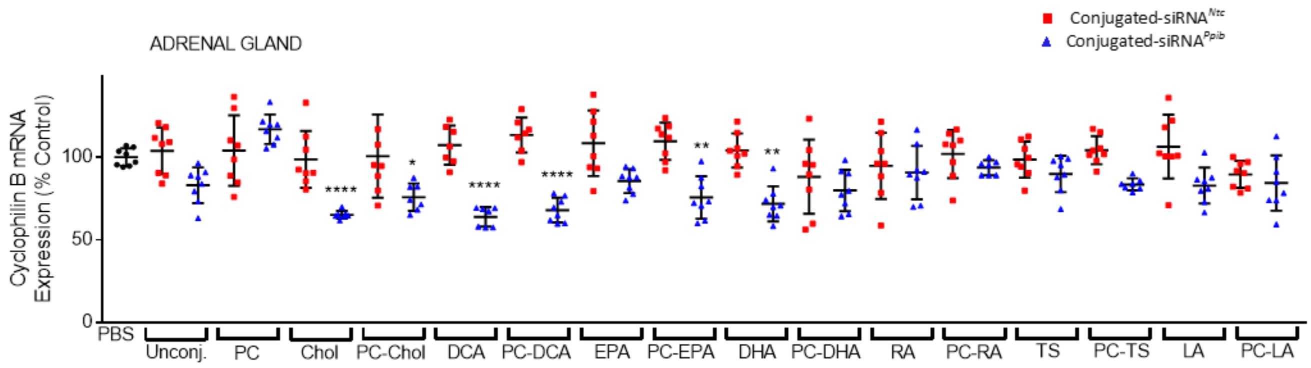


| | PBS | Unconj. | PC | Chol | PC-Chol | DCA | PC-DCA | EPA | PC-EPA | DHA | PC-DHA | RA | PC-RA | TS | PC-TS | LA | PC-LA |
|---|---|---|---|---|---|---|---|---|---|---|---|---|---|---|---|---|---|
| **Huntingtin mRNA silencing (% compared to *Ntc*)** | 0 ± 13 | 6 ± 15 | 20 ± 19 | 19 ± 10 | 14 ± 13 | 23 ± 12 | 47 ± 7 | 5 ± 12 | 12 ± 9 | 36 ± 23 | 36 ± 14 | 46 ± 16 | 10 ± 23 | 37 ± 9 | 21 ± 10 | 20 ± 17 | 19 ± 13 |
| **Significance (compared to *Ntc*)** | / | ns | ns | ns | ns | * | **** | ns | ns | * | * | **** | ns | ** | ns | ns | ns |
| **Cyclophilin B mRNA silencing (% compared to *Ntc*)** | 0 ± 5 | 21 ± 11 | 0 ± 9 | 34 ± 2 | 25 ± 8 | 46 ± 6 | 48 ± 7 | 23 ± 7 | 34 ± 13 | 32 ± 11 | 8 ± 13 | 4 ± 16 | 8 ± 5 | 9 ± 11 | 21 ± 4 | 24 ± 11 | 5 ± 17 |
| **Significance (compared to *Ntc*)** | / | ns | ns | ** | ns | **** | **** | ns | *** | ** | ns | ns | ns | ns | ns | ns | ns |

**Supplementary Figure 21: Efficacy of conjugated siRNAs in adrenal glands.** Subcutaneous injection (FVB/N mice); 20 mg/kg; collection of tissues one week after injection; n = 16 per gene and per conjugate (included non-targeting controls or *Ntc*). Huntingtin (*Htt*) (upper panel) and Cyclophilin B (*Ppib*) (lower panel) mRNA levels were measured using QuantiGene® (Affymetrix), normalized to a housekeeping gene, *Hprt* (Hypoxanthine-guanine phosphoribosyl transferase), and presented as percent of PBS (Phosphate buffered saline) control (mean ± SD). Data analysis: Outliners define with Grubb's method (alpha = 0.1%); Multiple comparisons = One-way ANOVA, Bonferroni test (****P<0.0001, ***P<0.001, **P<0.01, *P<0.1). The table indicates the average of inhibition percentages (n = 8) and significances for each target and conjugate compared to *Ntc* (mean ± SD; ns = non-significant).

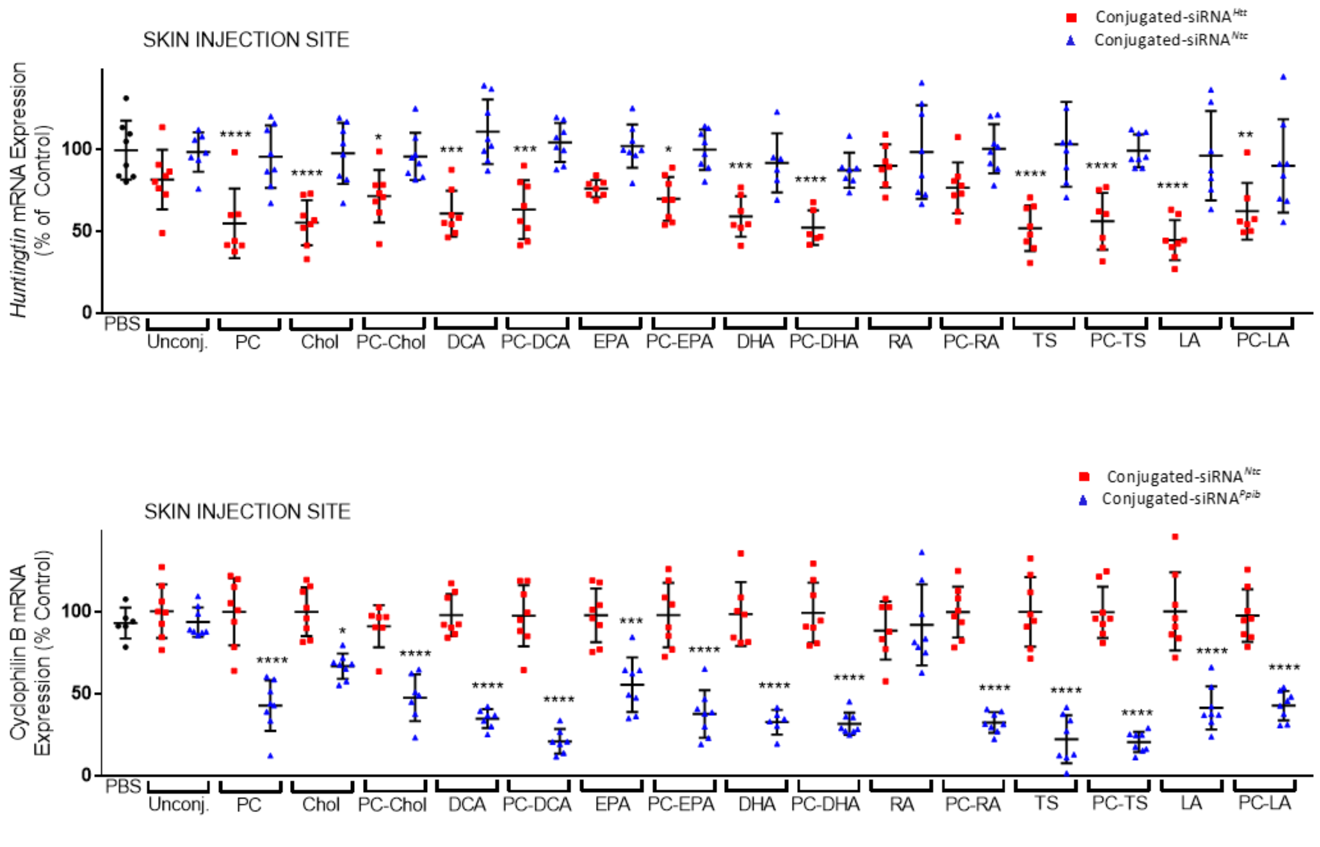


| | PBS | Unconj. | PC | Chol | PC-Chol | DCA | PC-DCA | EPA | PC-EPA | DHA | PC-DHA | RA | PC-RA | TS | PC-TS | LA | PC-LA |
|---|---|---|---|---|---|---|---|---|---|---|---|---|---|---|---|---|---|
| **Huntingtin mRNA silencing (% compared to *Ntc*)** | 0 ± 18 | 17 ± 18 | 41 ± 21 | 42 ± 14 | 24 ± 16 | 50 ± 14 | 41 ± 18 | 26 ± 5 | 30 ± 13 | 33 ± 12 | 35 ± 11 | 9 ± 13 | 24 ± 16 | 51 ± 14 | 43 ± 17 | 52 ± 12 | 28 ± 17 |
| **Significance (compared to *Ntc*)** | / | ns | ** | *** | * | **** | ** | ns | * | ** | ** | ns | ns | **** | *** | **** | ns |
| **Cyclophilin B mRNA silencing (% compared to *Ntc*)** | 0 ± 9 | 7 ± 9 | 58 ± 15 | 33 ± 8 | 44 ± 14 | 63 ± 6 | 77 ± 7 | 43 ± 17 | 60 ± 15 | 66 ± 8 | 68 ± 7 | 0 ± 25 | 67 ± 6 | 78 ± 15 | 79 ± 6 | 59 ± 13 | 55 ± 9 |
| **Significance (compared to *Ntc*)** | / | ns | **** | ** | *** | **** | **** | **** | **** | **** | **** | ns | **** | **** | **** | **** | **** |

**Supplementary Figure 22: Efficacy of conjugated siRNAs in skin (site of injection).** Subcutaneous injection (FVB/N mice); 20 mg/kg; collection of tissues one week after injection; n = 16 per gene and per conjugate (included non-targeting controls or *Ntc*). Huntingtin (*Htt*) (upper panel) and Cyclophilin B (*Ppib*) (lower panel) mRNA levels were measured using QuantiGene® (Affymetrix), normalized to a housekeeping gene, *Hprt* (Hypoxanthine-guanine phosphoribosyl transferase), and presented as percent of PBS (Phosphate buffered saline) control (mean ± SD). Data analysis: Outliners define with Grubb's method (alpha = 0.1%); Multiple comparisons = One-way ANOVA, Bonferroni test (****P<0.0001, ***P<0.001, **P<0.01, *P<0.1). The table indicates the average of inhibition percentages (n = 8) and significances for each target and conjugate compared to *Ntc* (mean ± SD; ns = non-significant).

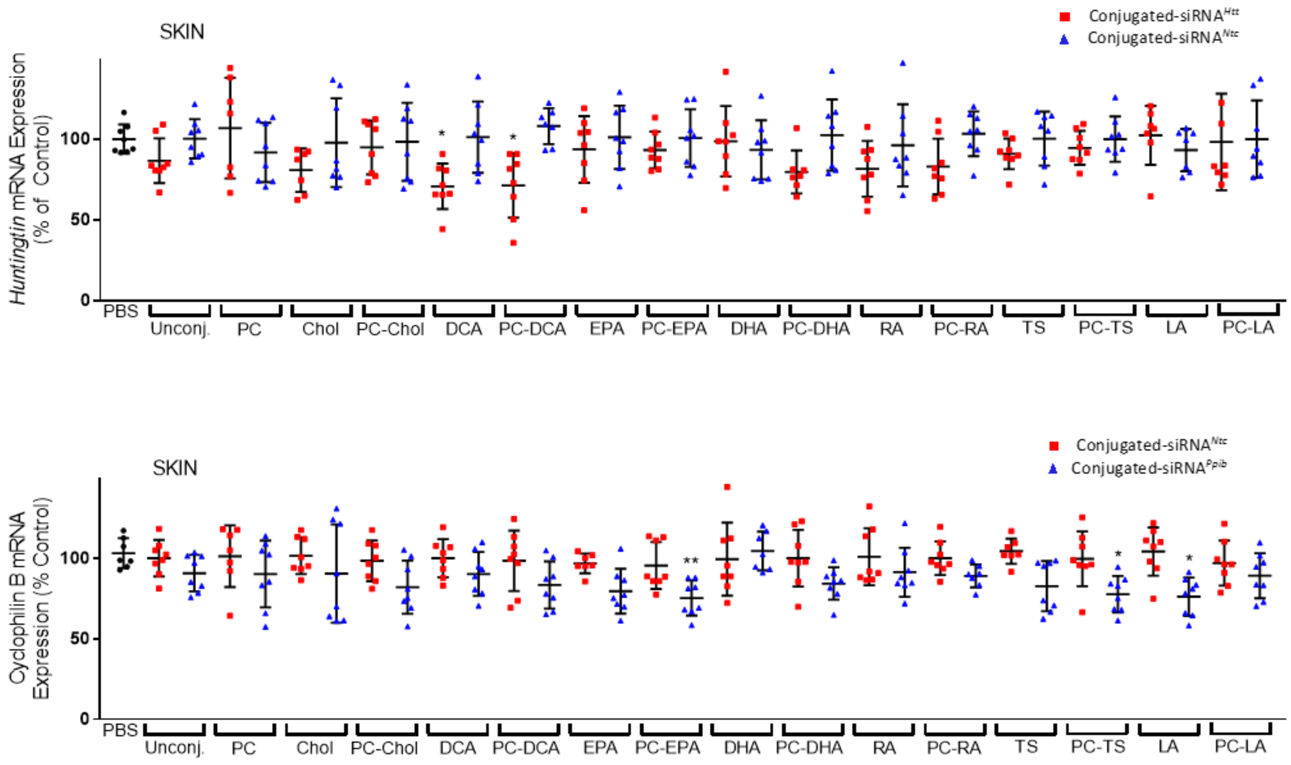


| | PBS | Unconj. | PC | Chol | PC-Chol | DCA | PC-DCA | EPA | PC-EPA | DHA | PC-DHA | RA | PC-RA | TS | PC-TS | LA | PC-LA |
|---|---|---|---|---|---|---|---|---|---|---|---|---|---|---|---|---|---|
| **Huntingtin mRNA silencing (% compared to *Ntc*)** | 0 ± 9 | 14 ± 14 | 0 ± 31 | 17 ± 13 | 4 ± 17 | 31 ± 14 | 37 ± 20 | 8 ± 21 | 7 ± 11 | 0 ± 22 | 23 ± 13 | 15 ± 17 | 20 ± 17 | 9 ± 10 | 5 ± 11 | 0 ± 18 | 2 ± 30 |
| **Significance (compared to *Ntc*)** | / | ns | ns | ns | ns | ns | * | ns | ns | ns | ns | ns | ns | ns | ns | ns | ns |
| **Cyclophilin B mRNA silencing (% compared to *Ntc*)** | 0 ± 9 | 9 ± 11 | 11 ± 21 | 11 ± 30 | 16 ± 17 | 10 ± 14 | 15 ± 15 | 27 ± 14 | 20 ± 11 | 0 ± 12 | 16 ± 10 | 10 ± 15 | 11 ± 7 | 22 ± 16 | 22 ± 11 | 28 ± 12 | 8 ± 14 |
| **Significance (compared to *Ntc*)** | / | ns | ns | ns | ns | ns | ns | ns | ns | ns | ns | ns | ns | ns | ns | ns | ns |

**Supplementary Figure 23: Efficacy of conjugated siRNAs in skin.** Subcutaneous injection (FVB/N mice); 20 mg/kg; collection of tissues one week after injection; n = 16 per gene and per conjugate (included non-targeting controls or *Ntc*). Huntingtin (*Htt*) (upper panel) and Cyclophilin B (*Ppib*) (lower panel) mRNA levels were measured using QuantiGene® (Affymetrix), normalized to a housekeeping gene, *Hprt* (Hypoxanthine-guanine phosphoribosyl transferase), and presented as percent of PBS (Phosphate buffered saline) control (mean ± SD). Data analysis: Outliners define with Grubb's method (alpha = 0.1%); Multiple comparisons = One-way ANOVA, Bonferroni test (**P<0.01, *P<0.1). The table indicates the average of inhibition percentages (n = 8) and significances for each target and conjugate compared to *Ntc* (mean ± SD; ns = non-significant).

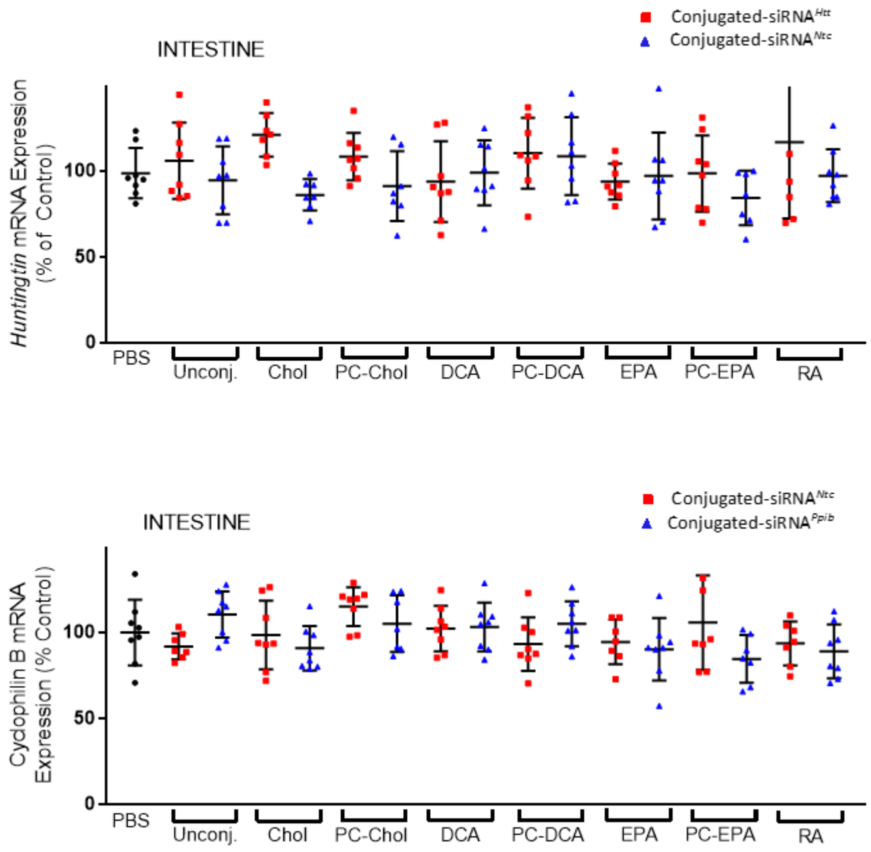


| | PBS | Unconj. | Chol | PC-Chol | DCA | PC-DCA | EPA | PC-EPA | RA |
|---|---|---|---|---|---|---|---|---|---|
| Huntingtin mRNA silencing (% compared to *Ntc*) | 0 ± 15 | 0 ± 22 | 0 ± 13 | 0 ± 14 | 5 ± 23 | 0 ± 21 | 3 ± 11 | 0 ± 22 | 0 ± 44 |
| Significance (compared to *Ntc*) | / | ns | ns | ns | ns | ns | ns | ns | ns |
| Cyclophilin B mRNA silencing (% compared to *Ntc*) | 0 ± 19 | 0 ± 13 | 8 ± 16 | 10 ± 13 | 0 ± 14 | 0 ± 13 | 4 ± 18 | 21 ± 14 | 5 ± 15 |
| Significance (compared to *Ntc*) | / | ns | ns | ns | ns | ns | ns | ns | ns |

**Supplementary Figure 24: Efficacy of conjugated siRNAs in intestine.** Subcutaneous injection (FVB/N mice); 20 mg/kg; collection of tissues one week after injection; n = 16 per gene and per conjugate (included non-targeting controls or *Ntc*). Huntingtin (*Htt*) (upper panel) and Cyclophilin B (*Ppib*) (lower panel) mRNA levels were measured using QuantiGene® (Affymetrix), normalized to a housekeeping gene, *Hprt* (Hypoxanthine-guanine phosphoribosyl transferase), and presented as percent of PBS (Phosphate buffered saline) control (mean ± SD). Data analysis: Outliners define with Grubb's method (alpha = 0.1%); Multiple comparisons = One-way ANOVA, Bonferroni test. The table indicates the average of inhibition percentages (n = 8) and significances for each target and conjugate as percent of *Ntc* (mean ± SD; ns = non-significant).

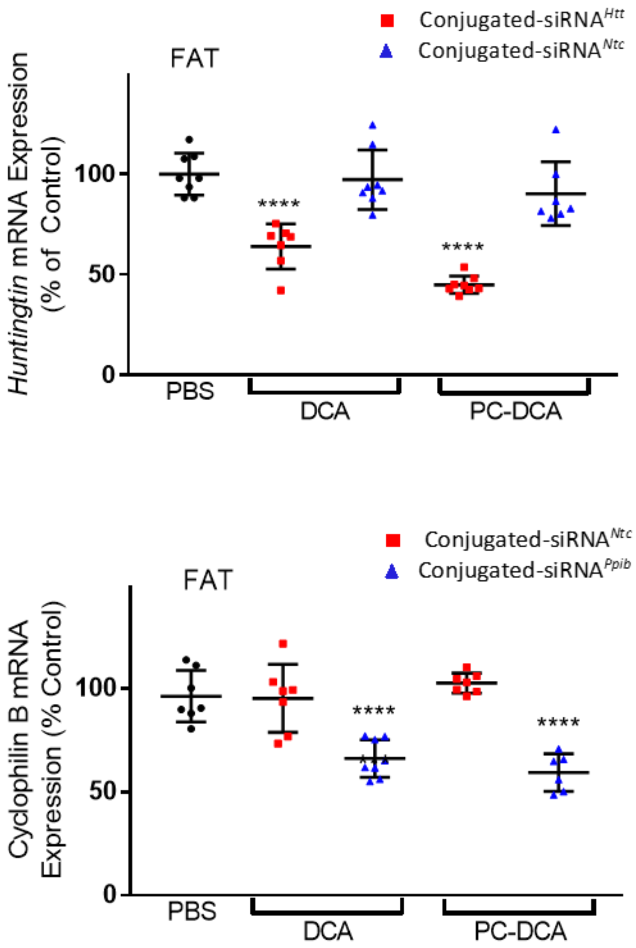


| | PBS | DCA | PC-DCA |
|---|---|---|---|
| **Huntingtin mRNA silencing (% compared to *Ntc*)** | 0 ± 10 | 30 ± 18 | 51 ± 4 |
| **Significance (compared to *Ntc*)** | / | **** | **** |
| **Cyclophilin B mRNA silencing (% compared to *Ntc*)** | 0 ± 16 | 35 ± 9 | 31 ± 20 |
| **Significance (compared to *Ntc*)** | / | *** | **** |

**Supplementary Figure 25: Efficacy of conjugated siRNAs in fat.** Subcutaneous injection (FVB/N mice); 20 mg/kg; collection of tissues one week after injection; n = 16 per gene and per conjugate (included non-targeting controls or *Ntc*). Huntingtin (*Htt*) (upper panel) and Cyclophilin B (*Ppib*) (lower panel) mRNA levels were measured using QuantiGene® (Affymetrix), normalized to a housekeeping gene, *Hprt* (Hypoxanthine-guanine phosphoribosyl transferase), and presented as percent of PBS (Phosphate buffered saline) control (mean ± SD). Data analysis: Outliners define with Grubb's method (alpha = 0.1%); Multiple comparisons = One-way ANOVA, Bonferroni test (****P<0.0001, ***P<0.001). The table indicates the average of inhibition percentages (n = 8) and significances for each target using DCA and PC-DCA conjugated siRNAs compared to *Ntc* (mean ± SD).

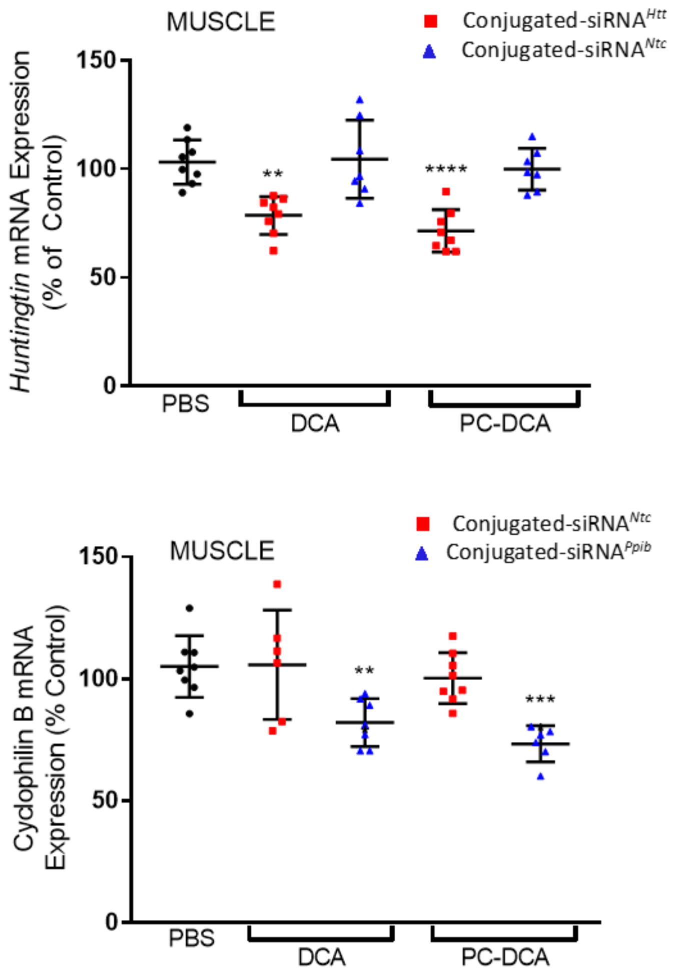


| | PBS | DCA | PC-DCA |
|---|---|---|---|
| Huntingtin mRNA silencing (% compared to *Ntc*) | 0 ± 10 | 21 ± 9 | 24 ± 10 |
| Significance (compared to *Ntc*) | / | ** | *** |
| Cyclophilin B mRNA silencing (% compared to *Ntc*) | 0 ± 13 | 29 ± 16 | 30 ± 11 |
| Significance (compared to *Ntc*) | / | ns | *** |

**Supplementary Figure 26: Efficacy of conjugated siRNAs in muscle.** Subcutaneous injection (FVB/N mice); 20 mg/kg; collection of tissues one week after injection; n = 16 per gene and per conjugate (included non-targeting controls or *Ntc*). Huntingtin (*Htt*) (upper panel) and Cyclophilin B (*Ppib*) (lower panel) mRNA levels were measured using QuantiGene® (Affymetrix), normalized to a housekeeping gene, *Hprt* (Hypoxanthine-guanine phosphoribosyl transferase), and presented as percent of PBS (Phosphate buffered saline) control (mean ± SD). Data analysis: Outliners define with Grubb's method (alpha = 0.1%); Multiple comparisons = One-way ANOVA, Bonferroni test (****P<0.0001, ***P<0.001, **P<0.01). The table indicates the average of inhibition percentages (n = 8) and significances for each target using DCA and PC-DCA conjugated siRNAs compared to *Ntc* (mean ± SD ; ns = non-significant).

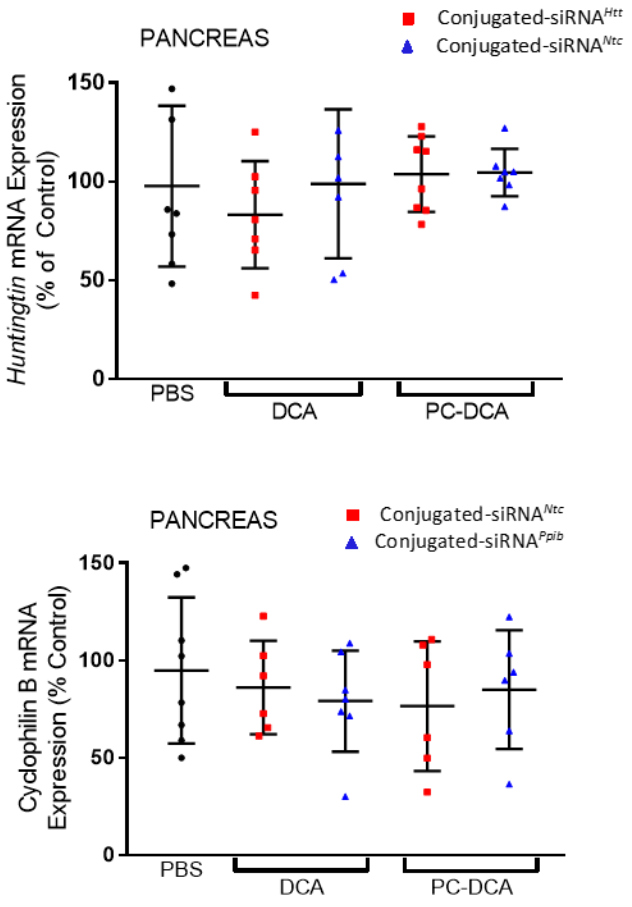


| | PBS | DCA | PC-DCA |
|---|---|---|---|
| Huntingtin mRNA silencing (% compared to *Ntc*) | 0 ± 41 | 16 ± 17 | 0 ± 19 |
| Significance (compared to *Ntc*) | / | ns | ns |
| Cyclophilin B mRNA silencing (% compared to *Ntc*) | 0 ± 37 | 7 ± 26 | 0 ± 30 |
| Significance (compared to *Ntc*) | / | ns | ns |

**Supplementary Figure 27: Efficacy of conjugated siRNAs in pancreas.** Subcutaneous injection (FVB/N mice); 20 mg/kg; collection of tissues one week after injection; n = 16 per gene and per conjugate (included non-targeting controls or *Ntc*). Huntingtin (*Htt*) (upper panel) and Cyclophilin B (*Ppib*) (lower panel) mRNA levels were measured using QuantiGene® (Affymetrix), normalized to a housekeeping gene, *Hprt* (Hypoxanthine-guanine phosphoribosyl transferase), and presented as percent of PBS (Phosphate buffered saline) control (mean ± SD). Data analysis: Outliners define with Grubb's method (alpha = 0.1%); Multiple comparisons = One-way ANOVA, Bonferroni test. The table indicates the average of inhibition percentages (n = 8) and significances for each target using DCA and PC-DCA conjugated siRNAs compared to *Ntc* (mean ± SD; ns = non-significant).

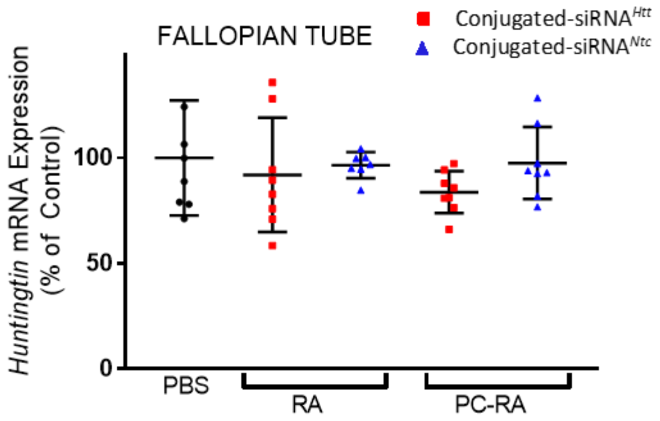


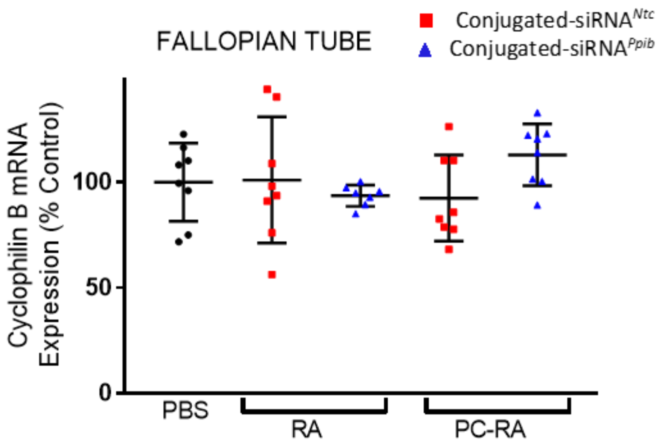


| | PBS | RA | PC-RA |
|---|---|---|---|
| Huntingtin mRNA silencing (% compared to *Ntc*) | 0 ± 27 | 5 ± 27 | 14 ± 10 |
| Significance (compared to *Ntc*) | / | ns | ns |
| Cyclophilin B mRNA silencing (% compared to *Ntc*) | 0 ± 18 | 7 ± 5 | 0 ± 15 |
| Significance (compared to *Ntc*) | / | ns | ns |

**Supplementary Figure 28: Efficacy of conjugated siRNAs in fallopian tube.** Subcutaneous injection (FVB/N mice); 20 mg/kg; collection of tissues one week after injection; n = 16 per gene and per conjugate (included non-targeting controls or *Ntc*). Huntingtin (*Htt*) (upper panel) and Cyclophilin B (*Ppib*) (lower panel) mRNA levels were measured using QuantiGene® (Affymetrix), normalized to a housekeeping gene, *Hprt* (Hypoxanthine-guanine phosphoribosyl transferase), and presented as percent of PBS (Phosphate buffered saline) control (mean ± SD). Data analysis: Outliners define with Grubb's method (alpha = 0.1%); Multiple comparisons = One-way ANOVA, Bonferroni test. The table indicates the average of inhibition percentages (n = 8) and significances for each target using RA and PC-RA conjugated siRNAs compared to *Ntc* (mean ± SD; ns = non-significant).

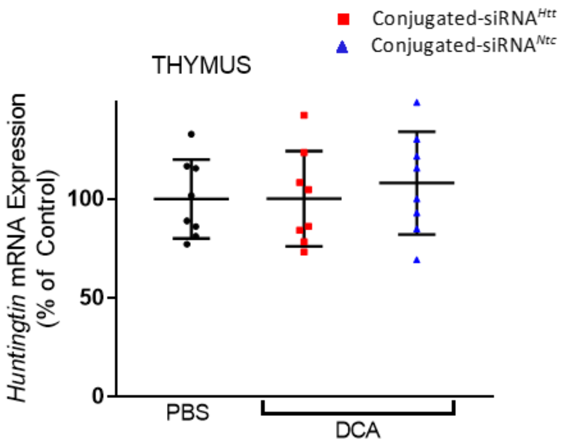


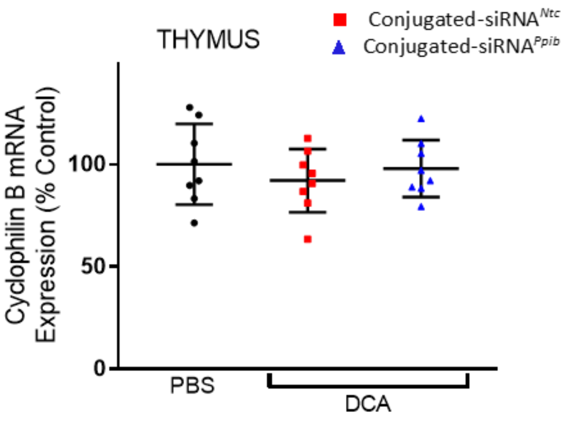


| | PBS | DCA |
|---|---|---|
| Huntingtin mRNA silencing (% compared to *Ntc*) | 0 ± 20 | 8 ± 24 |
| Significance (compared to *Ntc*) | / | ns |
| Cyclophilin B mRNA silencing (% compared to *Ntc*) | 0 ± 20 | 0 ± 14 |
| Significance (compared to *Ntc*) | / | ns |

**Supplementary Figure 29: Efficacy of conjugated siRNAs in thymus.** Subcutaneous injection (FVB/N mice); 20 mg/kg; collection of tissues one week after injection; n = 16 per gene and per conjugate (included non-targeting controls or *Ntc*). Huntingtin (*Htt*) (upper panel) and Cyclophilin B (*Ppib*) (lower panel) mRNA levels were measured using QuantiGene® (Affymetrix), normalized to a housekeeping gene, *Hprt* (Hypoxanthine-guanine phosphoribosyl transferase), and presented as percent of PBS (Phosphate buffered saline) control (mean ± SD). Data analysis: Outliners define with Grubb's method (alpha = 0.1%); Multiple comparisons = One-way ANOVA, Bonferroni test. The table indicates the average of inhibition percentages (n = 8) and significances for each target using DCA conjugated siRNAs compared to *Ntc* (mean ± SD; ns = non-significant).

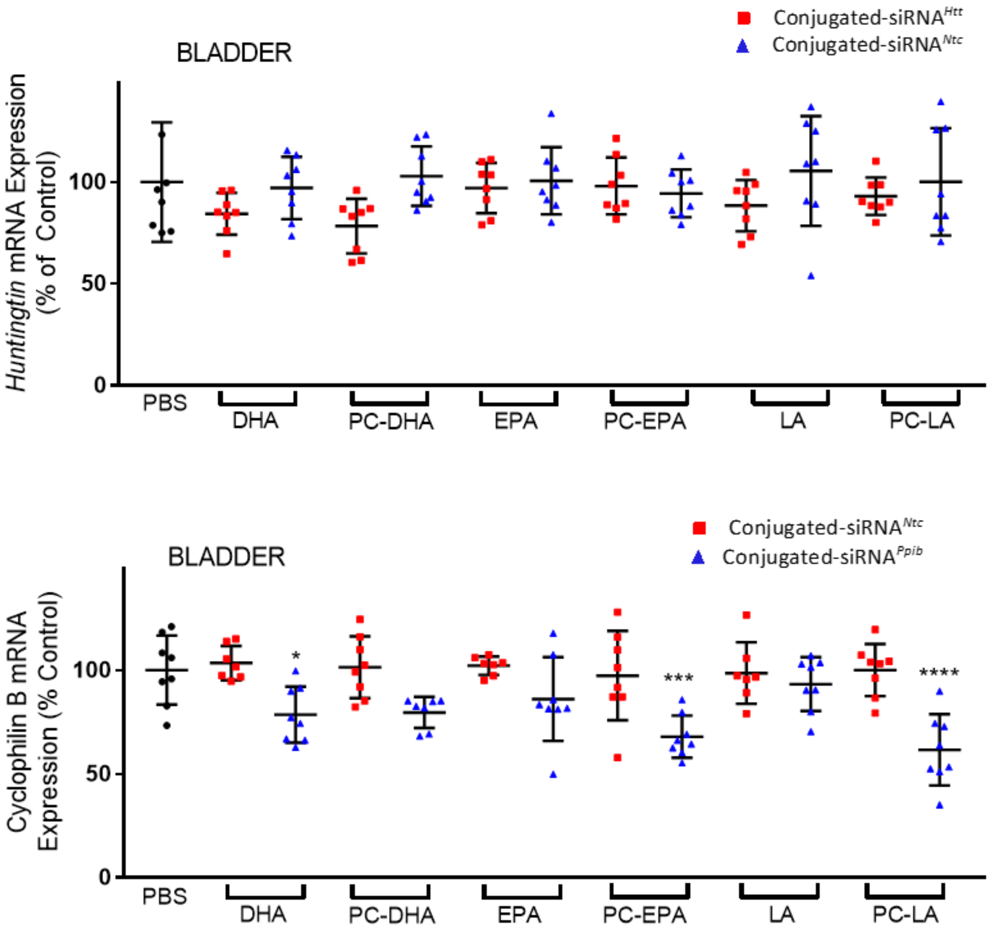


| | PBS | DHA | PC-DHA | EPA | PC-EPA | LA | PC-LA |
|---|---|---|---|---|---|---|---|
| Huntingtin mRNA silencing (% compared to *Ntc*) | 0 ± 29 | 13 ± 10 | 25 ± 13 | 4 ± 12 | 0 ± 14 | 17 ± 13 | 7 ± 9 |
| Significance (compared to *Ntc*) | / | ns | ns | ns | ns | ns | ns |
| Cyclophilin B mRNA silencing (% compared to *Ntc*) | 0 ± 17 | 25 ± 14 | 22 ± 8 | 16 ± 20 | 30 ± 10 | 5 ± 13 | 39 ± 17 |
| Significance (compared to *Ntc*) | / | * | ns | ** | * | ns | **** |

**Supplementary Figure 30: Efficacy of conjugated siRNAs in bladder.** Subcutaneous injection (FVB/N mice); 20 mg/kg; collection of tissues one week after injection; n = 16 per gene and per conjugate (included non-targeting controls or *Ntc*). Huntingtin (*Htt*) (upper panel) and Cyclophilin B (*Ppib*) (lower panel) mRNA levels were measured using QuantiGene® (Affymetrix), normalized to a housekeeping gene, *Hprt* (Hypoxanthine-guanine phosphoribosyl transferase), and presented as percent of PBS (Phosphate buffered saline) control (mean ± SD). Data analysis: Outliners define with Grubb's method (alpha = 0.1%); Multiple comparisons = One-way ANOVA, Bonferroni test (****P<0.0001, ***P<0.001, **P<0.01, *P<0.1). The table indicates the average of inhibition percentages (n = 8) and significances for each target and conjugates compared to *Ntc* (mean ± SD; ns = non-significant).

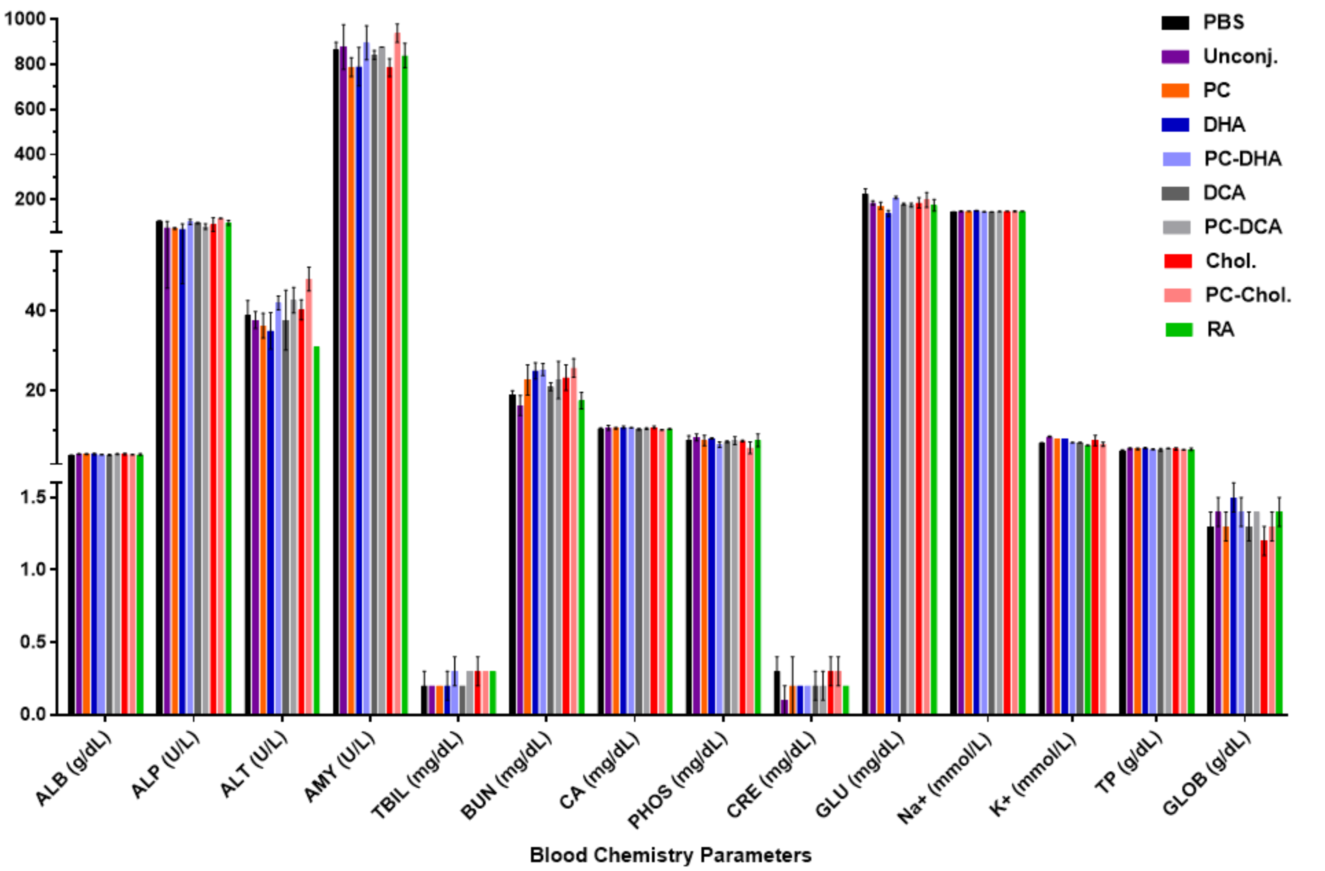


| | PBS | Unconj. | PC | DHA | PC-DHA | DCA | PC-DCA | Chol. | PC-Chol. | RA |
|---|---|---|---|---|---|---|---|---|---|---|
| ALB (g/dL) | 3.7 ± 0.2 | 4.1 ± 0.1 | 4.1 ± 0.1 | 4.1 ± 0.2 | 4.0 ± 0.1 | 3.9 ± 0.2 | 4.1 ± 0.1 | 4.1 ± 0.2 | 4.0 ± 0.1 | 4.0 ± 0.2 |
| ALP (U/L) | 105.3 ± 1.5 | 74.7 ± 29.0 | 72.0 ± 3.5 | 69.3 ± 22.5 | 101.0 ± 11.3 | 96.0 ± 3.0 | 80.3 ± 12.1 | 89.3 ± 30.1 | 116.7 ± 1.5 | 97.0 ± 11.3 |
| ALT (U/L) | 39.0 ± 3.6 | 37.7 ± 2.1 | 36.3 ± 3.1 | 35.0 ± 4.6 | 42.0 ± 1.7 | 37.7 ± 7.5 | 42.7 ± 3.2 | 40.3 ± 2.5 | 48.0 ± 3.0 | 31.0 ± 0.0 |
| AMY (U/L) | 864.3 ± 33.5 | 877.0 ± 99.0 | 788.0 ± 41.7 | 789.7 ± 85.2 | 895.7 ± 75.4 | 841.7 ± 19.5 | 876.6 ± 0.7 | 785.7 ± 39.5 | 938.0 ± 41.0 | 839.0 ± 53.7 |
| TBIL (mg/dL) | 0.2 ± 0.1 | 0.2 ± 0.0 | 0.2 ± 0.0 | 0.2 ± 0.1 | 0.3 ± 0.1 | 0.2 ± 0.0 | 0.3 ± 0.0 | 0.3 ± 0.1 | 0.3 ± 0.0 | 0.3 ± 0.0 |
| BUN (mg/dL) | 19.0 ± 1.0 | 16.3 ± 2.5 | 22.7 ± 3.8 | 25.0 ± 2.0 | 25.3 ± 1.5 | 21.0 ± 1.0 | 22.7 ± 4.7 | 23.3 ± 3.2 | 25.7 ± 2.3 | 17.5 ± 2.1 |
| CA (mg/dL) | 10.5 ± 0.2 | 10.7 ± 0.6 | 10.6 ± 0.2 | 10.8 ± 0.3 | 10.7 ± 0.1 | 10.3 ± 0.2 | 10.5 ± 0.2 | 10.8 ± 0.3 | 10.2 ± 0.1 | 10.4 ± 0.1 |
| PHOS (mg/dL) | 7.7 ± 1.0 | 8.3 ± 0.9 | 7.5 ± 1.3 | 8.0 ± 0.2 | 6.5 ± 0.7 | 7.2 ± 0.2 | 7.5 ± 1.0 | 7.4 ± 0.2 | 5.7 ± 1.5 | 7.6 ± 1.6 |
| CRE (mg/dL) | 0.3 ± 0.1 | 0.1 ± 0.1 | 0.2 ± 0.2 | 0.2 ± 0.2 | 0.2 ± 0.0 | 0.2 ± 0.0 | 0.2 ± 0.1 | 0.3 ± 0.1 | 0.3 ± 0.1 | 0.2 ± 0.0 |
| GLU (mg/dL) | 225.7 ± 22.4 | 185.7 ± 9.3 | 172.7 ± 15.3 | 138.3 ± 14.0 | 209.0 ± 6.2 | 180.0 ± 3.5 | 175.7 ± 8.1 | 185.3 ± 23.5 | 199.0 ± 33.2 | 175.0 ± 25.5 |
| Na+ (mmol/L) | 144.7 ± 1.5 | 148.3 ± 1.5 | 148.3 ± 0.6 | 150.0 ± 2.0 | 146.0 ± 1.0 | 145.3 ± 1.5 | 147.3 ± 0.6 | 148.0 ± 1.0 | 148.0 ± 2.0 | 148.5 ± 0.7 |
| K+ (mmol/L) | 6.9 ± 0.1 | 8.5 ± 0.1 | 7.9 ± 0.0 | 8.0 ± 0.0 | 7.0 ± 0.1 | 7.0 ± 0.1 | / | 7.5 ± 1.3 | 6.6 ± 0.5 | 6.3 ± 0.1 |
| TP (g/dL) | 5.0 ± 0.2 | 5.5 ± 0.2 | 5.4 ± 0.2 | 5.6 ± 0.2 | 5.3 ± 0.1 | 5.2 ± 0.3 | 5.5 ± 0.1 | 5.4 ± 0.3 | 5.2 ± 0.1 | 5.3 ± 0.3 |
| GLOB (g/dL) | 1.3 ± 0.1 | 1.4 ± 0.1 | 1.3 ± 0.1 | 1.5 ± 0.1 | 1.4 ± 0.1 | 1.3 ± 0.1 | 1.4 ± 0.0 | 1.2 ± 0.1 | 1.3 ± 0.1 | 1.4 ± 0.1 |

**Supplementary Figure 31. Blood chemistry parameters for conjugated siRNAs treated mice.** Subcutaneous injection (FVB/N mice); 20 mg/kg; collection of blood one week after injection; n = 3 per group. ALB = Albumin; ALP = Alkaline Phosphatase; ALT = Alanine Aminotransferase; AMY = Amylase; TBIL = Total Bilirubin; BUN = Blood Urea Nitrogen; CA = Calcium; PHOS = Phosphate; CRE = Creatinine; GLU = Glucose; Na+ = Sodium; K+ = Potassium; TP = Total Protein; GLOB = Globulin.
